# Quantitative analyses reveal extracellular dynamics of Wnt ligands in *Xenopus* embryos

**DOI:** 10.1101/2020.02.20.957860

**Authors:** Yusuke Mii, Kenichi Nakazato, Chan-Gi Pack, Yasushi Sako, Atsushi Mochizuki, Shinji Takada, Masanori Taira

**Affiliations:** National Institute for Basic Biology and Exploratory Research Center on Life and Living Systems (ExCELLS), National Institutes of Natural Sciences, 5-1 Higashiyama, Myodaiji, Okazaki, Aichi 444-8787, Japan; The Graduate University for Advanced Studies (SOKENDAI), Okazaki, Aichi 444-8787, Japan; JST, PRESTO, 4-1-8 Honcho, Kawaguchi, Saitama, 332-0012, Japan; Department of Biological Sciences, Graduate School of Science, University of Tokyo, 7-3-1 Hongo, Bunkyo-ku, Tokyo 113-0033, Japan; Theoretical Biology Laboratory, RIKEN, 2-1 Hirosawa, Wako 351-0198, Japan; Department of Computational Intelligence and Systems Science, Interdisciplinary Graduate School of Science and Engineering, Tokyo Institute of Technology, Yokohama 226-8503, Japan; Cellular Informatics Laboratory, RIKEN, 2-1 Hirosawa, Wako 351-0198, Japan

**Keywords:** diffusion, morphogen, Wnt, sFRP, heparan sulfate, FCS, FRAP, FDAP

## Abstract

The mechanism of intercellular transport of Wnt ligands is still a matter of debate. Here, to better understand this issue, we examined distribution and dynamics of Wnt8 in *Xenopus* embryos. While Venus-tagged Wnt8 was found on the surfaces of cells close to Wnt-producing cells, we also detected its dispersal over distances of 15 cell diameters. A combination of fluorescence correlation spectroscopy and quantitative imaging revealed that only a small proportion of Wnt8 ligands diffuses freely, whereas most Wnt8 molecules are bound to cell surfaces. Fluorescence decay after photoconversion showed that Wnt8 ligands bound on cell surfaces decreased exponentially, suggesting a dynamic exchange of bound forms of Wnt ligands. Mathematical modelling based on this exchange recapitulates a graded distribution of bound, but not free, Wnt ligands. Based on these results, we propose that Wnt distribution in tissues is controlled by a dynamic exchange of its abundant bound and rare free populations.

## INTRODUCTION

The Wnt family of secreted signaling proteins has versatile roles in animal development, stem cell systems, and carcinogenesis (Clevers et al., 2014; Loh et al., 2016; Nusse and Clevers, 2017). It has been generally accepted that in the extracellular space, morphogenic Wnt ligands form a concentration gradient by dispersal (Clevers et al., 2014; Kiecker and Niehrs, 2001; Muller et al., 2013; Smith, 2009; Strigini and Cohen, 2000; Tabata and Takei, 2004; Yan and Lin, 2009; Zecca et al., 1996; Zhu and Scott, 2004). In contrast to this classical view, evidence also suggests dispersal-independent functions of Wnt ligands. For instance, a membrane-tethered form of Wingless (Wg) can recapitulate an almost normal pattern of *Drosophila* wings, suggesting that dispersal of Wg is dispensable for patterning (Alexandre et al., 2014). This dispersal-independent patterning can be explained by gradual attenuation of Wg expression in distally localized cells in which Wg was formerly expressed. However, it remains unclear to what extent dispersal-dependent and/or -independent mechanisms contribute to the graded distribution of Wnt proteins in tissue patterning.

Visualization of Wnt ligands is essential to understand their distributions. In the wing disc of *Drosophila*, Wg proteins are widely distributed from wing margin cells, where Wg is expressed (Strigini and Cohen, 2000; Zecca et al., 1996). Furthermore, long-range dispersal of Wg was evidenced by an experiment in which Wg was captured by distally expressed Frizzled2, a Wg receptor (Chaudhary et al., 2019). Similarly, endogenous Wnt ligands tagged with fluorescent proteins showed long-range distributions in *C. elegans* (Pani and Goldstein, 2018). In addition to these observations in invertebrates, we found that endogenous Wnt8 ligands disperse far from their source cells in *Xenopus* embryos (Mii et al., 2017). On the other hand, mouse Wnt3 accumulates within a few cell diameters of its source cells in the microenvironment of the intestine (Farin et al., 2016). These studies show that Wnt ligands apparently disperse in tissues and embryos, although the dispersal range varies. Importantly, in many of these studies, Wnt ligands accumulate locally on cell surfaces, showing punctate distributional patterns (Pani and Goldstein, 2018; Strigini and Cohen, 2000; Zecca et al., 1996). Furthermore, we demonstrated that Wnt8 and Frzb, a secreted Wnt inhibitor, accumulate separately and locally on cell surfaces in *Xenopus* embryos (Mii et al., 2017). However, these punctate accumulations on cell surfaces, largely ignored in the literature, in the context of Wnt gradient formation, raise the question of whether such accumulations contribute to formation of concentration gradients in tissues and embryos.

Studies in *Drosophila* wing disc have shown that cell surface scaffolds, such as heparan sulfated (HS) proteoglycans (HSPGs), are required for both distribution and delivery of morphogens, including Wg, Hedgehog (Hh), and Decapentaplegic (Dpp) (Franch-Marro et al., 2005; Lin, 2004; Yan and Lin, 2009). From these studies, the “restricted diffusion” model, in which morphogens are transferred extracellularly by interacting with cell surface scaffolds, has been proposed (Yan and Lin, 2009). In this model, the movement of each morphogen molecule is constrained in a “bucket brigade” fashion by interactions with cell surface scaffolds. As a result of continuous interactions, morphogen molecules are slowly transferred (Han et al., 2005, Yan, 2009 #152; Kerszberg and Wolpert, 1998; Takei et al., 2004). However, it is difficult to explain local accumulations of Wnt proteins by the restricted diffusion mechanism, because passive diffusion alone should result in smoothly decreasing gradients. However, we recently showed that HSPGs on cell surfaces are discretely distributed in a punctate manner, which varies with heparan sulfate (HS) modification, forming two different types of HS clusters, *N*-sulfo-rich and *N*-acetyl-rich forms (Mii et al., 2017). Notably, Wnt8 and Frzb, a secreted Frizzled-related protein (sFRP), accumulate separately on *N-*sulfo-rich and *N-*acetyl-rich HS clusters, respectively. Frzb expands the distribution and signaling range of Wnt8 by forming heterocomplexes (Mii and Taira, 2009), and Wnt8/Frzb complexes are colocalized with *N-*acetyl-rich HS clusters (Mii et al., 2017). *N*-sulfo-rich clusters are frequently internalized together with Wnt8, whereas *N*-acetyl-rich HS clusters tend to remain on the cell surface. This difference in stability on the cell surface may account for the short-range distribution of Wnt8 and the long-range distribution of Frzb (Mii and Taira, 2009; Mii et al., 2017) and suggests that the distribution of HS clusters should be considered in order to understand extracellular dynamics of Wnt ligands.

To explain the dynamics of Wnt ligands in tissues, quantitative analyses of Wnt ligands are required. Dynamics of secreted proteins have been investigated using fluorescence recovery after photobleaching (FRAP) (Sprague and McNally, 2005; Sprague et al., 2004) and fluorescence correlation spectroscopy (FCS) (Hess et al., 2002; Kicheva et al., 2012; Muller et al., 2013). For example, FRAP measurements have shown that Dpp and Wg diffuse slowly in the *Drosophila* wing disc with diffusion coefficients ranging from 0.05 to 0.10 μm^2^/s, suggestive of the restricted diffusion model (Kicheva et al., 2007). In contrast, FCS measurements of FGF8 in zebrafish embryos showed fast, virtually free diffusion, with a diffusion coefficient of ∼50 μm^2^/s (Yu et al., 2009). Furthermore, in contrast to the FRAP results, free diffusion of Dpp measured in the *Drosophila* wing disc using FCS yielded a diffusion coefficient of ∼20 μm^2^/s (Zhou et al., 2012). Thus, diffusion coefficients measured with FRAP and FCS differ by 3-4 orders of magnitude (Rogers and Schier, 2011). FCS is based on fixed-point scanning within a confocal volume (typically sub-femtoliter) for several seconds, while FRAP evaluates considerably larger regions of photobleaching/photoconversion, containing tens or hundreds of cells (Rogers and Schier, 2011) and spanning long time windows (typically several hours). Under these experimental conditions for FRAP, it is proposed that diffusion of secreted proteins is affected by zigzag paths of the narrow intercellular space between polygonal epithelial cells, instead of an open, unobstructed space (hindered diffusion model) (Muller et al., 2013), and/or by endocytosis, which reduces the concentration of the diffusing species in the extracellular space. Thus, FRAP data should not be directly compared with data obtained by FCS analysis.

In this study, we examined extracellular dynamics of Wnt8 and Frzb, both of which are involved in anteroposterior patterning of vertebrate embryos (Clevers and Nusse, 2012; Kiecker and Niehrs, 2001; MacDonald et al., 2009; Mii et al., 2017). First, we visualized their localization in *Xenopus* embryos by fusing them with fluorescent proteins and we examined their dispersion by capturing them in distant cells. We also examined their dispersal dynamics using FCS- and fluorescence decay after photoconversion (FDAP) measurements in embryonic tissue. In particular, we refined FDAP-based analysis by focusing on a limited area at the cell boundary, which enabled us to quantify dynamics comparable to those measured by FCS. Based on these results and our previous findings, we propose a basic mathematical model to explain distribution and dispersion of secreted proteins.

## RESULTS

### Extracellular distributions of secreted proteins depend on interactions with molecules on cell surfaces

As we have previously shown, Wnt8 and Frzb fused with monomeric Venus (mV) were visualized on cell surfaces when expressed in *Xenopus* embryos (Mii et al., 2017). In contrast, we found that only the secreted form of mV (sec-mV), which was not expected to bind specifically to the cell surface, was hardly visible along the cell boundary under the same conditions (Figure 1A, right). Since Wnt8 and Frzb colocalize with heparan sulfate clusters on cell surfaces, we speculated that binding to cell surface proteins, like heparan sulfate proteoglycans (HSPGs), affects the distribution of Wnt8 and Frzb. To examine this possibility, we added heparin-binding (HB) peptides, consisting of 16 (ARKKAAKA)_2_ (HB2) or 32 amino acids (ARKKAAKA)_4_ (HB4) (Verrecchio et al., 2000) (Figure 1C) to sec-mV. Addition of HB peptides significantly increased the intensity of mVenus fluorescence in the intercellular region compared to that of sec-mV. This suggests that the intercellular distribution of secreted proteins depends on interactions with docking molecules on cell surfaces.

**Figure 1.**
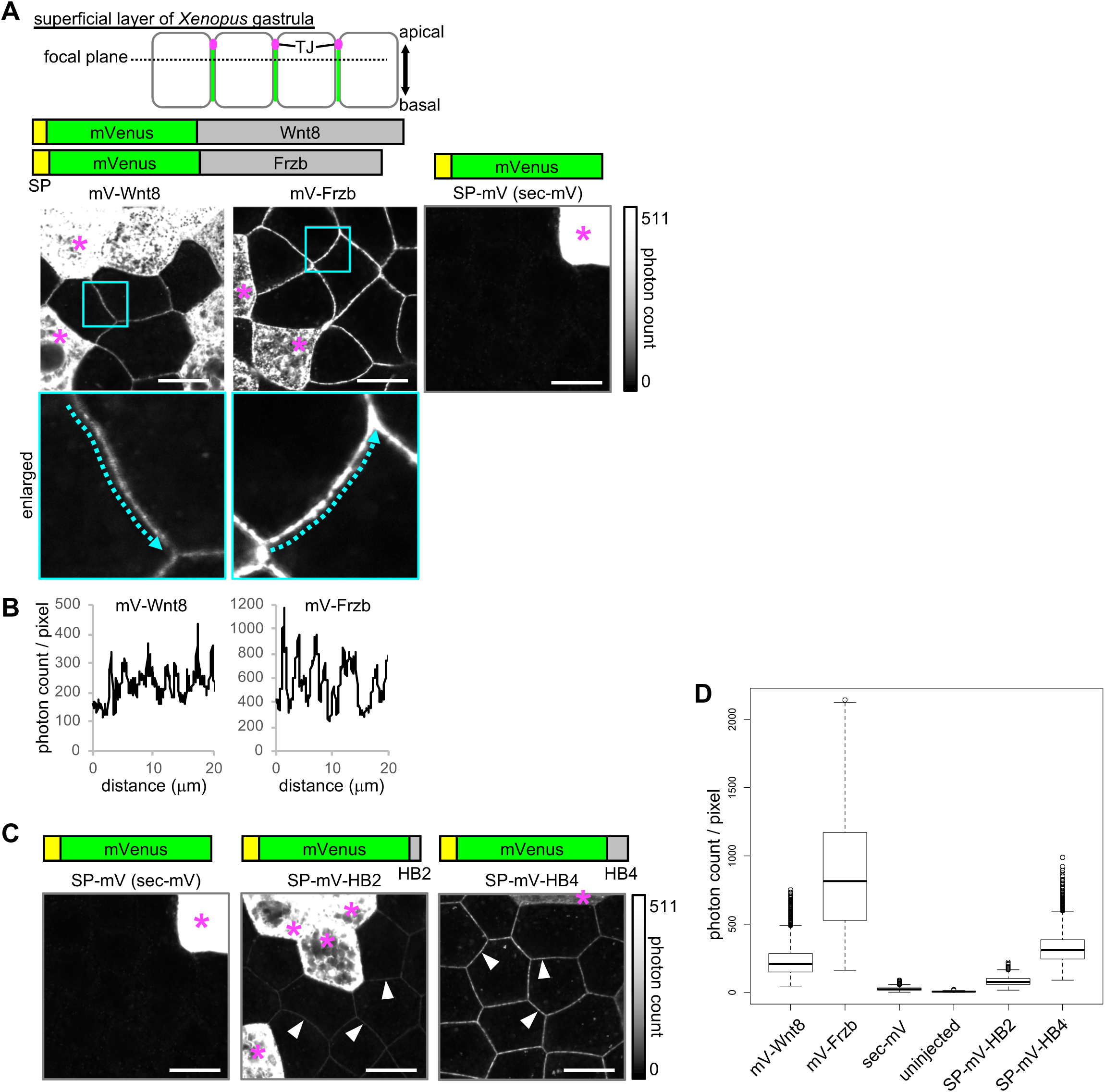
Extracellular distribution of Wnt8, Frzb, and artificial secreted proteins. All images presented were acquired using live-imaging with photon counting method, which enables saturation-free imaging even with samples with a wide dynamic range. (**A**) Distribution of secreted proteins in the superficial layer of living *Xenopus* gastrula (st. 10.5-11.5). Observed focal planes were at subapical level as illustrated. mRNAs for indicated mVenus (mV) fusion proteins were microinjected into a single ventral blastomere of 4- or 8-cell stage embryos to observe regions adjacent to the source cells (indicated with asterisks). All images were acquired in the same condition with photon counting detection. Look up tables (LUT) show the range of the photon count in the images. (B) Intensity plots for mV-Wnt8 and mV-Frzb in the intercellular space. Plots along the arrows in enlarged pictures in (A) are shown. (C) Distribution of artificial secreted proteins in *Xenopus* embryos. The data of sec-mV is the same as in (A). sec-mV was not apparent in the intercellular space, whereas sec-mV-HB2 and sec-mV-HB4 were distributed in the intercellular space (arrowheads). SP, signal peptide; HB, heparin binding peptide. (D) Quantification of fluorescent intensities in the intercellular space. Photon count per pixel are presented. All samples show statistically significant difference to each other (*p* < 2e-16, pairwise comparisons using Wilcoxon rank sum test). Scale bars, 20 μm. Amounts of injected mRNAs (ng/embryo): *mV-wnt8*, *mV-frzb*, *sp-mV*, *sp-mV-hb2*, or *sp-mV-hb4*, 0.25.

To directly examine this idea, we constructed a reconstitution system, consisting of HA-epitope-tagged secreted mVenus (sec-mV-2HA) and a membrane-tethered anti-HA antibody (“tethered-anti-HA Ab”) (Figure 2A, see Figure 2-Figure supplement 1 for cDNA cloning and validation of anti-HA antibody). This artificial protein and tethered-anti-HA Ab were expressed in separated areas in the animal cap region of *Xenopus* gastrulae. As with sec-mV, sec-mV-2HA was hardly visible in the intercellular space, even close to the source cells (Figure 2B). In contrast, sec-mV-2HA was observed around tethered-anti-HA Ab-expressing cells that were traced with memRFP, even though these cells were distantly located from the source cells (Figure 2B). Thus, interaction with cell surface proteins can affect distributions of secreted proteins.

**Figure 2.**
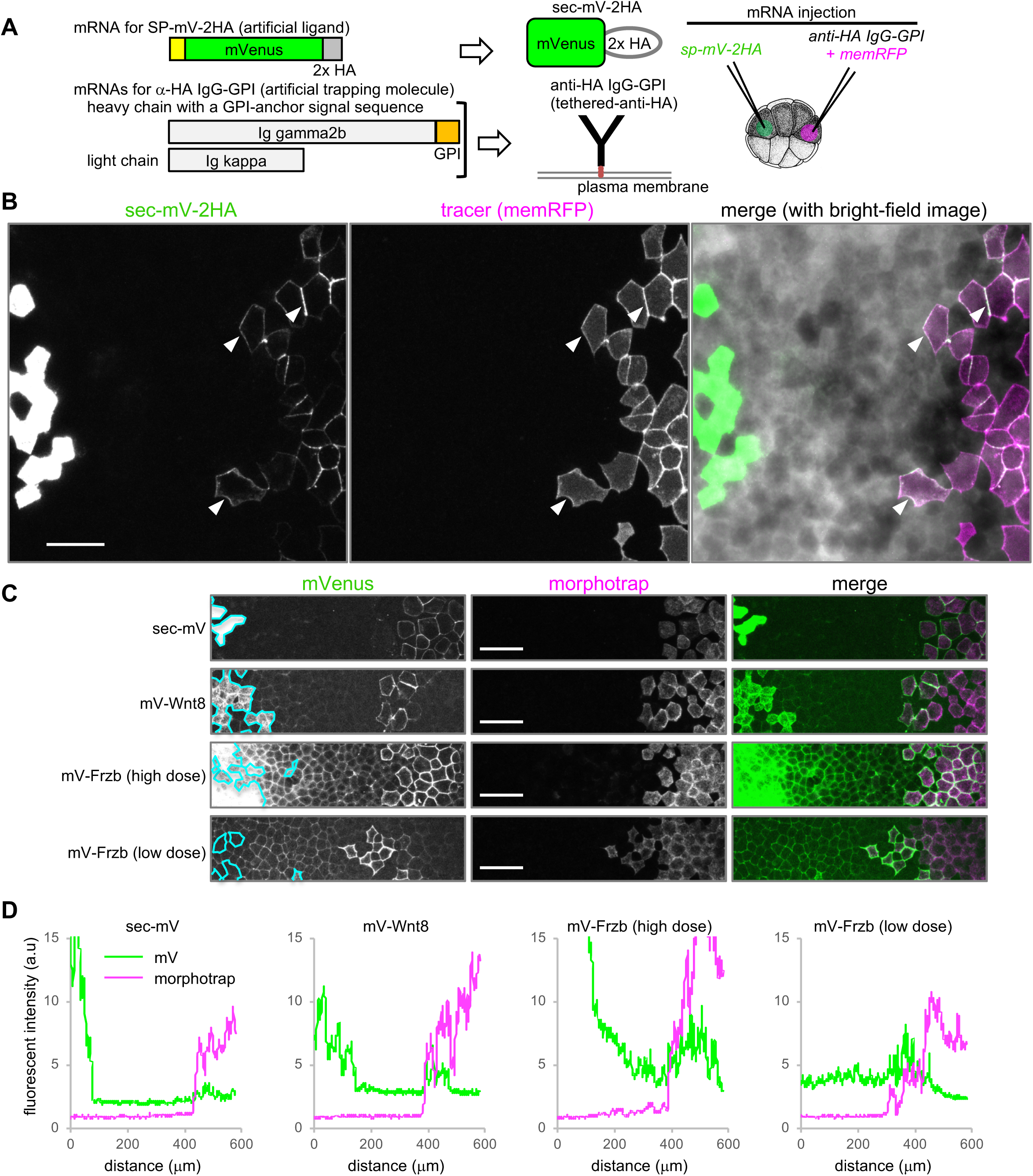
tethered-anti-HA Ab and morphotrap. (A) Schematic representation of tethered-anti-HA Ab. (B) Result of tethered-anti-HA Ab. The artificial ligand (sec-mV-2HA) was trapped at the tethered-anti-HA Ab expressing cells at a distant region from the source. Superficial layer of *Xenopus* gastrula (st. 11.5) was imaged as a z-stack and its maximum intensity projection (MIP) was presented for the fluorescent images. Intercellular mVenus signal (green) of sec-mV-2HA was not apparent in the vicinity of source cells, but was detected memRFP (magenta). (C) Morphotrap at a distant region from the source. Superficial layer of *Xenopus* gastrula (st. 11.5) was imaged as a z-stack and its maximum intensity projection (MIP) was presented for the fluorescent images. Intercellular mVenus signal of an artificial ligand, sec-mV (green), was not detected in the vicinity of source cells (green) (left panel), but was detected around the morphotrap-expressing cells that can be traced by mCherry fluorescence (middle panels). Also mV-Wnt8 and mV-Frzb were trapped and accumulated on the distant morphotrap-expressing cells, suggesting the existence of diffusing molecules in the distant region. The source regions are indicated with cyan lines according to memBFP (tracer for mV-tagged proteins, not shown). (D) Distribution of mVenus and morphotrap. Fluorescent intensity of mVenus and mCherry (for morphotrap) was plotted from the left to the right. Scale bars, 100 μm. Amounts of injected mRNAs (ng/embryo) *sp-mV-2ha*, 1.0; *memRFP*, 0.15; *ig gamma2b-gpi*, 1.1; *ig kappa*, 0.63 (B); *sec-mV, mV-wnt8*, or *mV-frzb* (high dose), 0.25; *mV-frzb* (low dose), 0.063; *morphotrap*, 1.0; *memBFP,* 0.1 (C).

This result also indicates that diffusing proteins are not readily visible using standard confocal microscopy, unless they are trapped by cell surface proteins. In fact, quantitative analysis of artificial secreted proteins revealed a slight, but significant, increase of photon counts in the intercellular region by injection of mRNA for *sec-mV*, compared to uninjected embryos, indicating that sec-mV actually exists in the intercellular region (Figure 1D, Figure 1-Figure supplement 1).

### Populations of secreted Wnt8 and Frzb proteins disperses long distance

We next examined dispersal of molecules of mV-Wnt8 and mV-Frzb. Both mV-Wnt8 and mV-Frzb accumulated locally along the cell boundary at the subapical level (Figure 1A and B), consistent with previous observations (Mii et al., 2017), which indicated that populations of Wnt8 and Frzb in the intercellular space were bound to the cell surface at HS clusters. On the other hand, given that some molecules of mV-Wnt8 or mV-Frzb may drift away from the cell surface, these proteins would be almost undetectable with standard confocal microscopy, as exemplified by sec-mV (Figure 1A) and tethered-anti-HA Ab (Figure 2B). To examine such mobile proteins, we tried to capture them using a “morphotrap” located distantly from the source cells (Figure 2C). A morphotrap is a membrane-tethered form of anti-GFP nanobody, originally devised to block dispersal of Dpp-GFP from source cells (Harmansa et al., 2015). We supposed that morphotraps could be utilized to detect or visualize diffusible proteins, similar to tethered-anti-HA Ab. As expected, sec-mV accumulated on the surface of morphotrap-expressing cells remote from source cells (Figure 2C). Similarly, mV-Wnt8 and mV-Frzb were trapped (Figure 2C), evidencing the long-distance (over 15 cells/200 μm) dispersal of some molecules. In addition to this dispersing population, mV-Wnt8 and mV-Frzb were also detectable in gradients from producing cells to morphotrap-expressing cells, unlike the case of sec-mV (Figure 2C and D). These results show that populations of mV-tagged Wnt8 and Frzb appeared not to associate tightly with cell surfaces, thereby potentially dispersing far from source cells.

### FCS analyses combined with quantitative imaging reveal cell-surface-bound and diffusing Wnt8 and Frzb proteins in the extracellular space

We next attempted to quantify the populations of Wnt8 or Frzb proteins associated with cell surfaces and diffusing in the extracellular space. For this purpose, we employed fluorescence correlation spectroscopy (FCS) to measure dynamics of mV-Wnt8, mV-Frzb, and sec-mV. FCS analyzes fluctuation of fluorescence by Brownian motion of fluorescent molecules in a sub-femtoliter confocal detection volume (Figure 3A-D). By autocorrelation analysis, FCS can measure the diffusion coefficient (*D*) and the number of particles, which is equivalent to the concentration of the diffusing molecules (Figure 3E), but it cannot measure immobile molecules (Hess et al., 2002).

**Figure 3.**
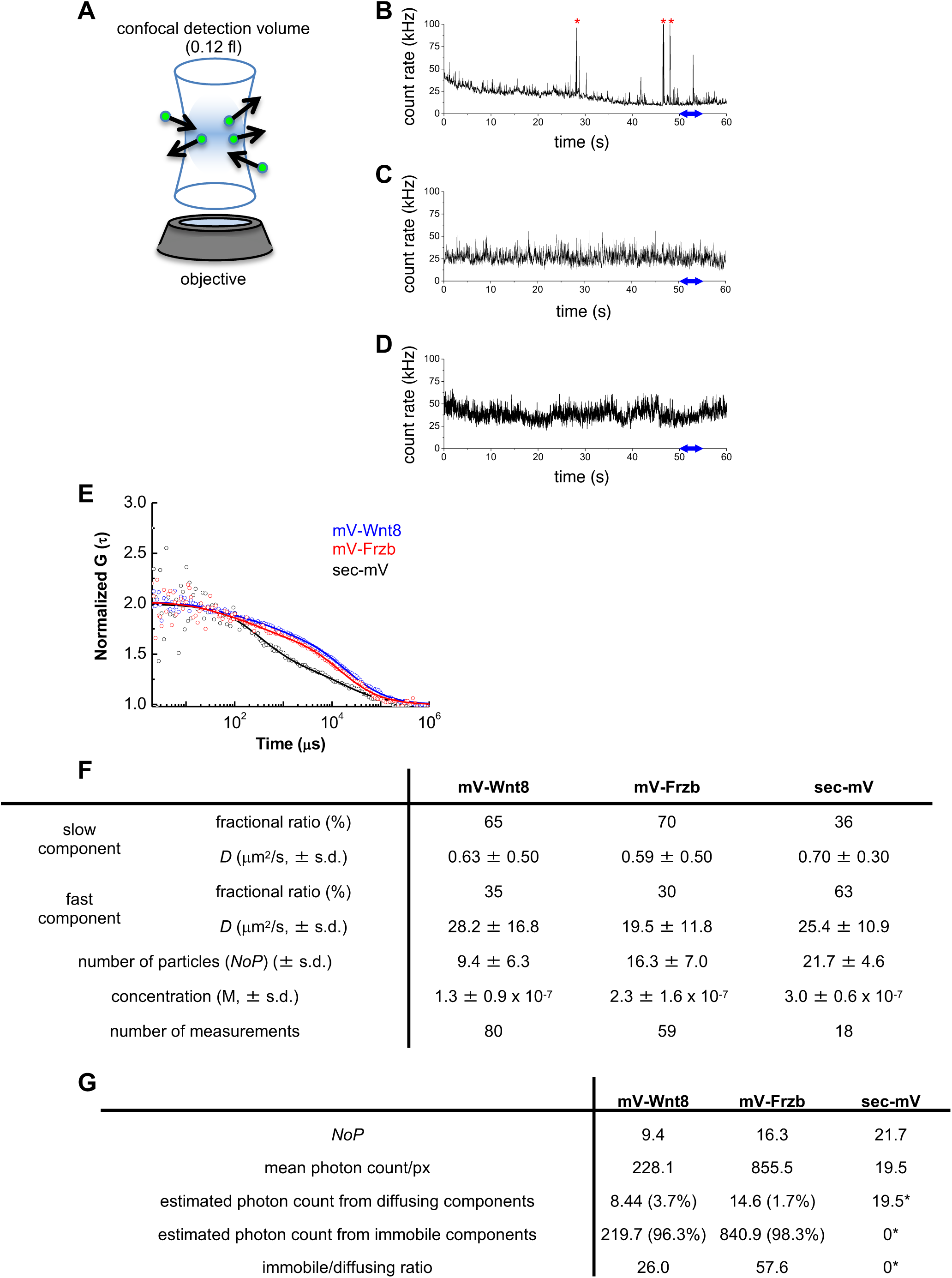
Fluorescence correlation spectroscopy (FCS) in the extracellular space of *Xenopus* embryos. mRNAs for mV-tagged proteins or sec-mV were injected into the animal pole region of a ventral blastomere of 4- or 8-cell stage *Xenopus* embryo. Injected embryos were observed at gastrula stages (st. 10.5-11.5). Each FCS measurement was performed at a point in the intercellular region within three cell diameters from the source cells. (A) Schematic illustration of FCS measurement. In FCS measurements, fluorescent signal usually fluctuates due to Brownian motion of the fluorescent molecules. Such fluctuations contain dynamic properties of the fluorescent molecules. Briefly, temporal frequency of the fluctuations corresponds to the diffusion coefficient (*D*) and amplitude of the fluctuations reversely corresponds to the number of particles in the confocal detection volume. (B-D) Time course of fluorescent intensities for mV-Wnt8 (B), mV-Frzb (C), and sec-mV (D), respectively. mV-Wnt8 shows characteristic peaks (asterisks). See also Figure supplement 1A-C for magnified views in a 5 seconds window as indicated by blue double headed arrows. (E) Autocorrelation functions of mV-Wnt8, mV-Frzb, and sec-mV. Data of mV-Wnt8, mV-Frzb, and sec-mV are plotted with blue, red, and black circles, respectively, with the best fitting curve. (F) Summary of FCS measurements. s.d., standard deviation. (G) Estimated proportion of diffusing and immobile components of mV-Wnt8 and mV-Frzb. Estimation is based on the assumption that sec-mV has no immobilized components (asterisks indicate assumed values). Amounts of injected mRNAs (ng/embryo): mV-wnt8, mV-frzb, and sec-mV, 0.25.

Using FCS, we previously showed that GFP-Wnt3a forms high-molecular-weight complexes in *Xenopus* embryos (Takada et al., 2018). Similarly, mV-Wnt8 shows characteristic peaks (Figure 3B, asterisks), suggesting formation of high-molecular-weight complexes of Wnt proteins. FCS measurements of these proteins were fitted to a two-component model, comprising fast and slow components. In the fast components, diffusion coefficients (*D*) of mV-Wnt8 and mV-Frzb were 28.2 ± 16.8 and 19.5 ± 11.8 µm^2^/s, respectively, comparable to that of sec-mV (25.4 ±10.9 µm^2^/s, mean ± s.d., Figures 3F and Figure 3-Figure supplement 1E). Values of *D_fast_* can be regarded as free diffusion, because theoretical, as well as reported *D* values of freely diffusing proteins of similar size, range from 10 to 100 µm^2^/s (Pack et al., 2006; Yu et al., 2009; Zhou et al., 2012). Importantly, diffusion coefficients of fast components of mV-Wnt8 and mV-Frzb were not significantly different from that of sec-mV, although the average percentage of the fast component was higher in sec-mV (63%) than mV-Wnt8 (35%) and mV-Frzb (30%) (Figures 3F and Figure 3-Figure supplement 1E). Thus, within the small volume of FCS measurements, a population of mV-Wnt8 and mV-Frzb molecules diffuse as rapidly as sec-mV molecules.

Importantly, we found that the number of particles (within the confocal volume, 0.12 fL), which is equivalent to the concentration of molecules measured by FCS, was somewhat lower in mV-Wnt8 (9.4 ± 6.3) and mV-Frzb (16.3 ± 7.0) than in sec-mV (21.7 ± 4.6) (Figure 3F and Figure 3—Figure supplement 1F). Because FCS analysis cannot detect immobile molecules (Hess et al., 2002), this result indicates that the concentration of diffusing molecules was relatively lower in mV-Wnt8 and mV-Frzb than in sec-mV. In contrast to this FCS result, quantitative analysis using confocal microscopy with photon counting imaging (reflecting both diffusing and immobile molecules) revealed that the amounts of mV-Wnt8 and mV-Frzb in the intercellular space was significantly higher than that of sec-mV (Figure 1D). Taken together, these results suggest that the percentage of immobilized mV-Wnt8 and mV-Frzb was much higher than of diffusing molecules measurable with FCS. We estimated the abundance of diffusing components as maximal, under the assumption that sec-mV has no immobile component (Figure 3G, assumed values are indicated with asterisks). Accordingly, the ratios of immobile to diffusing molecules were 26.0 for mV-Wnt8 and 57.6 for mV-Frzb (Figure 3G). These results suggest that only several percent of these proteins are diffusing, while the rest are bound to cell surfaces.

### FDAP analyses suggest exchange of cell-surface-bound and unbound states of Wnt8 and Frzb proteins

Although FCS analysis is suitable for measuring diffusing molecules, it cannot directly analyze molecules with extremely low mobility (Hess et al., 2002). To directly analyze dynamics of such molecules, we next employed fluorescence decay after photostimulation/photoconversion (FDAP) assays (Matsuda et al., 2008; Muller et al., 2012) in the intercellular space of *Xenopus* embryos.

Since FRAP/FDAP measurements usually examine considerably wider regions (typically containing tens or hundreds of cells) than with FCS (Rogers and Schier, 2011), direct comparisons of dynamics between FRAP/FDAP and FCS may not be relevant. Therefore, we restricted the area of photoconversion to a width of 1.66 μm and reduced the measurement time (16 sec), allowing us to obtain dynamic data in the intercellular region under conditions comparable to those for FCS (Figure 4A). We refer to this FDAP mode as “cell-boundary FDAP.” In this analysis, we fused a photoconvertible fluorescent protein, mKikGR (mK) (Habuchi et al., 2008) to Wnt8 and Frzb (mK-Wnt8 and mK-Frzb). These fusion proteins showed distributions in embryos similar to mV-tagged proteins (Figure 4B), and still retained biological activity (Figure 4-Figure supplement 1). Importantly, observed distribution patterns of mK-Wnt8 and mK-Frzb (Figure 4A) were quite stable for up to tens of minutes (data not shown). Therefore, we assumed a steady state during the FDAP analysis (16 sec).

**Figure 4.**
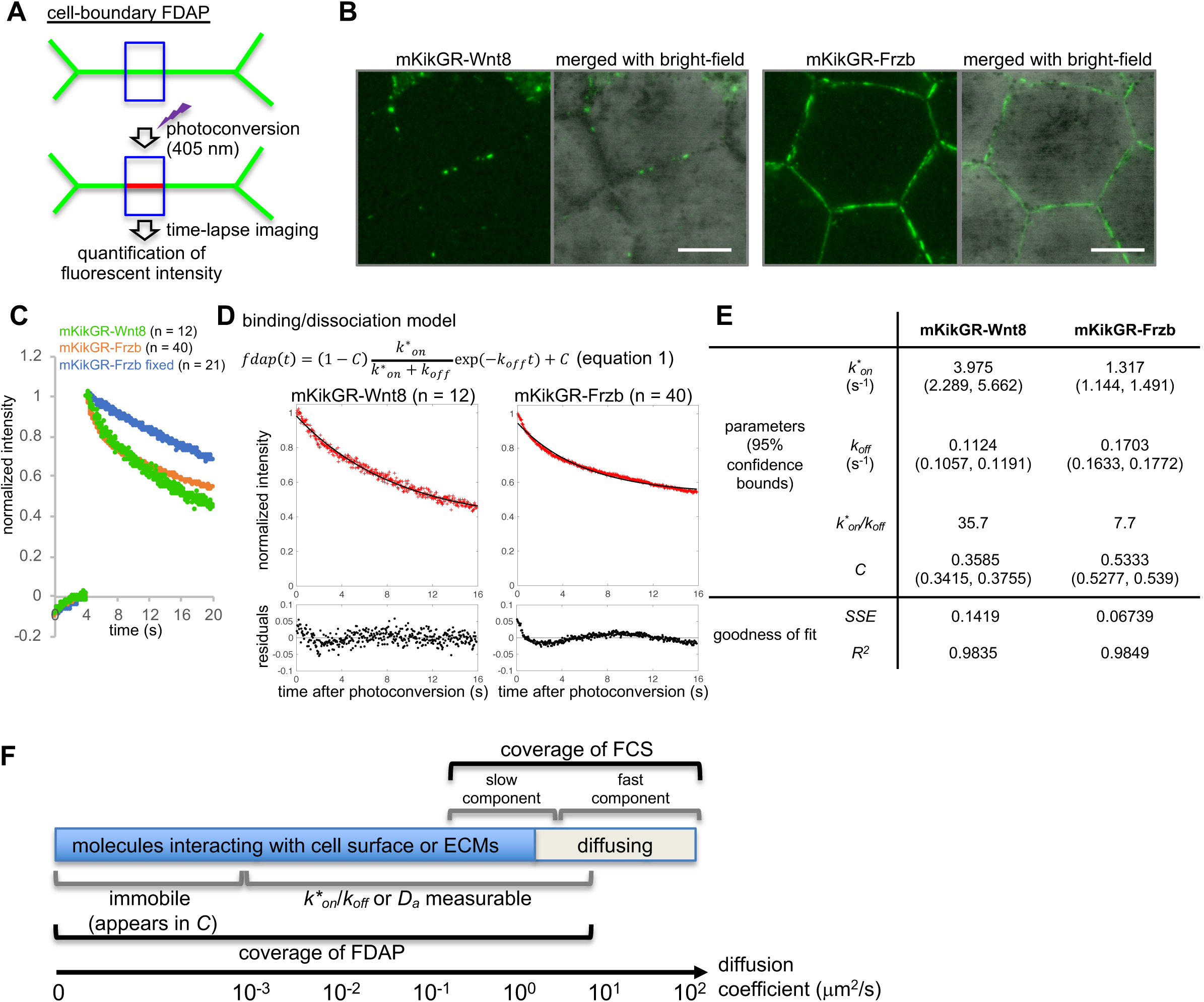
Fluorescence decay after photoconversion (FDAP) assay at the cell-boundary of Xenopus embryos. (A) Schematic illustration of cell-boundary FDAP assay. Green lines represent mKikGR-fusion protein distributed in the intercellular region. See also Videos 1-3 and the text for detail. (B) Intercellular distribution of mKikGR-Wnt8 and mKikGR-Frzb. Superficial layer of *Xenopus* gastrula (st. 10.5) was imaged as a z-stack and maximum intensity projection (MIP) was presented. Scale bars, 20 μm. (C) Time course of red (photoconverted state) fluorescent intensity within the photoconverted region. Photoconversion was performed about 4 seconds after the beginning of the measurement. Mean of normalized intensities were presented (for s.d., see Figure supplement 2A). Data of “mKikGR-Frzb fixed” were measured with MEMFA-fixed mKikGR-Frzb expressing embryos as immobilized controls. Numbers of measurements were indicated as n, which were collected in multiple experiments (twice for mKikGR-Wnt8 and mKikGR-Frzb fixed, and four times for mKikGR-Frzb). (D) Fluorescent decay curve fitted with the dissociation model. Mean of the normalized intensities for each time point was plotted with red cross. Fitting curves are shown as black lines. The residuals were mostly within 5% (0.05) and within 10% (0.1) in all cases. (E) Coefficients and evaluation of goodness of fit with the dissociation model. *k*_on_*, pseudo-on rate constant; *k_off_*, off-rate constant; *k*_on_/k_off_*, pseudo-equilibrium binding constant; *SSE*, sum of squared errors; *R*^2^, coefficient of determination. (F) Relationship of FCS-measurable and FDAP measurable components along with diffusion coefficient. Amounts of injected mRNAs (ng/embryo): *mkikGR-wnt8* and *mkikGR-frzb, 4.0*.

After photoconversion, red fluorescent intensity of mK-tagged proteins was measured within the same rectangular area as photoconversion (Figure 4A). Because puncta of Wnt8 are often internalized with HS clusters (Mii et al., 2017), we excluded data in which vesicular incorporation was observed during measurement of mK-Wnt8. As an immobilized control, mK-Frzb in formaldehyde-fixed embryos was similarly photoconverted. Its intensity within the rectangular region was fitted to the photobleaching model using repeated laser scanning, confirming that it actually was immobilized by formaldehyde fixation (Figure 4-Figure supplement 2B). Compared with this fixed control, mK-Wnt8 and mK-Frzb showed faster decline of fluorescent intensities (Figure 4C and Figure 4-Figure supplement 2A), confirming that abundance of mK-Wnt8 and mK-Frzb diminished in the photoconverted area.

FDAP spatial intensity profiles documented reduced abundance of photoconverted proteins over time in this area (Figure 4-Figure supplement 2C). However, photoconverted mK-Wnt8 did not accumulate in adjacent areas (Figure 4-Figure supplement 2C, 1st and 2nd panels, see also Videos 1 and 2). Probably, this is because diffusion of photoconverted proteins is so rapid that they cannot accumulate in this vicinity. This result suggests that photoconverted proteins did not move laterally, but instead moved far from the photoconverted area. On the other hand, mK-Frzb shows slight, but proximal accumulation, but this shift caused only a slight change in the spatial intensity profile (Figure 4-Figure supplement 2C, 3rd and 4th panels). Therefore, we conclude that these results closely resemble “diffusion-uncoupled FRAP/FDAP”, also known as a “reaction-dominant” case (Sprague and McNally, 2005; Sprague et al., 2004), in which pure diffusion is fast enough for molecules to diffuse away before binding in the vicinity of their origin. Indeed, FDAP curves of mK-Wnt8 and mK-Frzb were well fitted to the binding/dissociation model, which considers exchange of cell-surface-bound and unbound proteins (Figure 4D, equation 1; residuals was mostly within 5% and all within 10%) with indicated parameters (Figure 4E, the pseudo-on rate constant *k*_on_*, the off-rate constant *k_off_*, and the rate of the constantly bound component *C*; for individual data plot, see Figure 4-Figure supplement 2D). Based upon parameters obtained from averaged data, we estimated a pseudo-equilibrium binding constant (*k*_on_/k_off_*), equivalent to the ratio of bound/unbound components (35.4 for mK-Wnt8 and 7.7 for mK-Frzb) (Figure 4E). In addition, individual *k*\**_on_*/*k_off_* values for mK-Wnt8 and mK-Frzb, which did not differ significantly, range roughly from 2.5 to 20 (Figure 4—Figure supplement 2D). Thus, we conclude that most mK-Wnt8 and mK-Frzb molecules are bound, but can be exchanged with unbound molecules. We note that the pseudo-equilibrium binding constant may be underestimated because curve fitting does not consider photobleaching, and that curve fitting involves the constantly bound component, *C*, which indicates slower moving or immobile components during the elapsed experimental time (Figure 4F). Nevertheless, it is noteworthy that the ratio of bound to unbound components of mK-Wnt8, estimated by FDAP analysis (35.4-fold difference), was similar to that calculated from the combination of our quantitative imaging and FCS analysis of mV-Wnt8 (26.0-fold difference, Figure 3G). Taken together, all of the quantitative analyses performed in this study indicate that most Wnt8 and Frzb molecules are bound to cell surfaces, whereas only a small percentage is freely diffusing.

### Mathematically modeling diffusion and distribution of secreted proteins

Based on our quantitative imaging (Figure 1) of tethered-anti-HA Ab and morphotrap (Figure 2) FCS (Figure 3) and FDAP (Figure 4), we concluded that most Wnt8 and Frzb molecules are bound to cell surfaces, while small numbers of freely diffusing molecules exist in the extracellular space. Furthermore, we have already shown that Wnt8 and Frzb utilize different types of HS clusters, N-sulfo-rich and N-acetyl-rich, as cell-surface scaffolds, respectively (Mii et al., 2017). Thus, we examined whether free diffusion and binding to HS clusters can explain the extracellular distribution or gradient formation of secreted proteins, using mathematical modeling.

Here, we consider two states of ligands: free and bound. This model includes five dynamic processes: (i) ligand production, (ii) diffusion of free molecules in the intercellular space, (iii) binding of ligands to HS clusters on cell surfaces, (iv) release of bound molecules from HS clusters and (v) internalization of bound molecules into cells. In one-dimensional space, the model is written as:

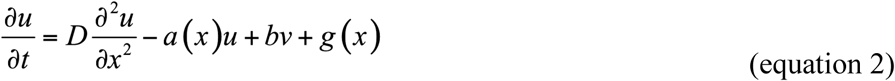

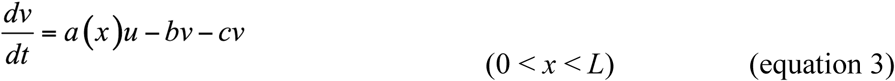

where *u* and *v* represent the concentration of free molecules and numbers of bound molecules, respectively, of a secreted protein. The symbols *x* and *t* are position and time, respectively. The symbols *a*(*x*), *b*, *c*, and *g*(*x*) represent binding, release, internalization, and production rates, respectively (Figure 5A); *a*(*x*) is equivalent to the amount of HS in HS clusters (for details, see Materials and Methods). *D* represents the diffusion coefficient of the free component in the extracellular space, which corresponds to diffusing components measured by FCS (Figure 3F).

**Figure 5.**
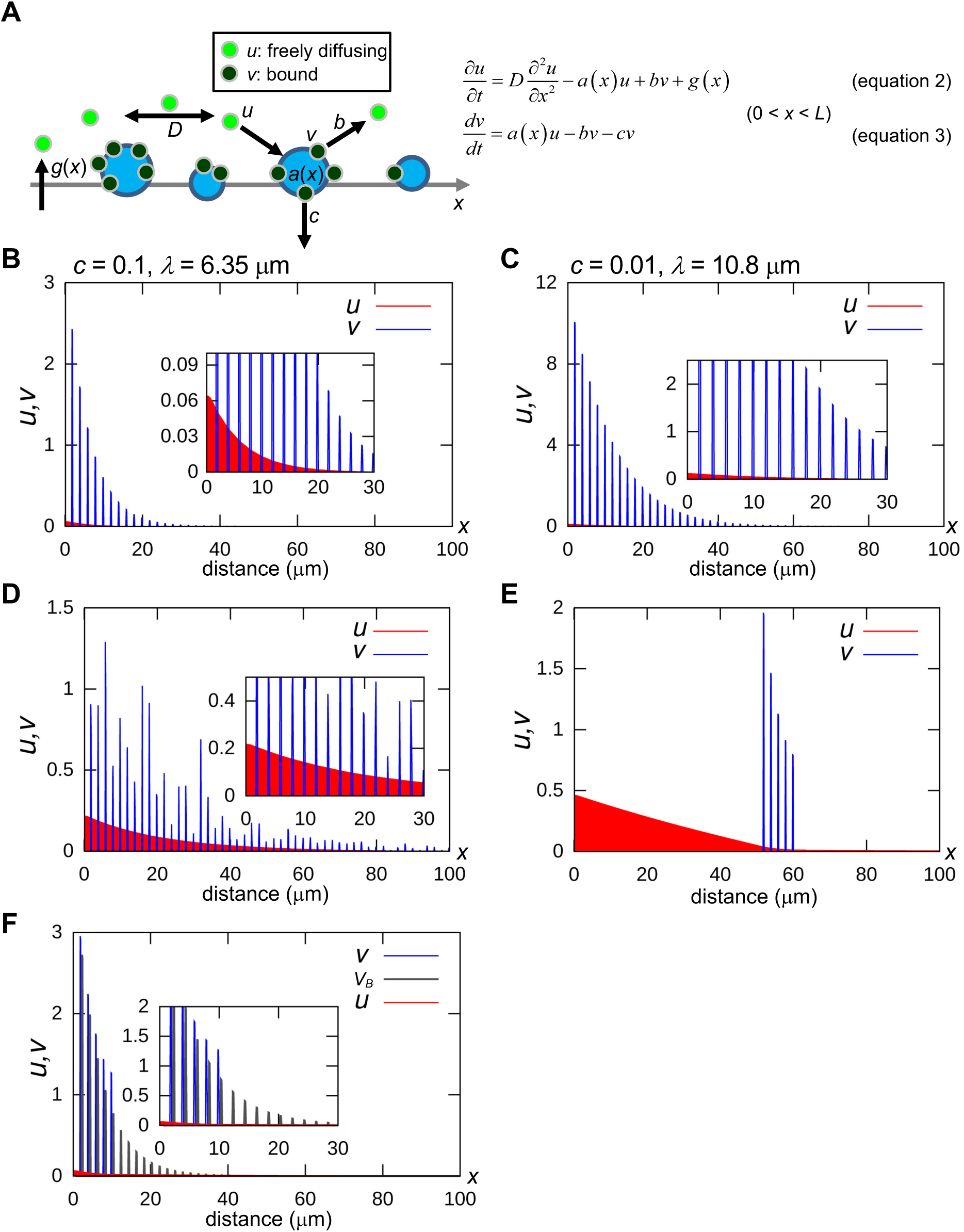
A minimal model of secreted protein dynamics in the extracellular space. Distributions of free (*u*) and bound (*v*) components of secreted proteins were obtained by computer simulation. The vertical axis indicates the amount of *u* and *v*, and the horizontal axis indicates the distance (*x*), where the spatial length *L* = 100 (μm). The distributions of *u* (red) and bound *v* (blue) at time *t* = 100 (sec) are shown, which we confirmed to be almost steady states. We used forward difference method with spatial step Δ*x* = 0.1 and temporal step Δ*t* = 0.0001 in numerical calculations. The level of *v* at the position where the docking sites exist (*a*(*x*) = *a*_n,max_) keeps relatively high even after enough time passed when *a*_n,max_ is higher than *b*. (A) Schema of the modeling. *a*(*x*), binding rate at position *x*. Note that *a*(*x*) is equivalent to the amount of HS for an HS cluster. *b*, release rate from the HS clusters. *c*, internalization rate of the HS clusters. *D*, diffusion coefficient of *u*. *g*(*x*), production rate at position *x*. For details, see Metarials and Methods. (B) Rapid internalization of the docking sites. Parameter values are: *D* = 20.0 (μm^2^/s), *a*_n,max_ = 10.0, *b* = 0.1, *c* = 0.1, *g*_max_ = 0.2, *R* = *L*/100, *p*_1_ = 2, and *p*_2_ = 0.2. The decay length *λ* is calculated as 6.35 μm according to the fitting curve (see Figure5-Figure supplement 1B). (C) Slow internalization of the docking sites. Parameter values are the same as in (A) except for *c* = 0.01. The decay length *λ* is calculated as 10.8 μm according to the fitting curve (see Figure5-Figure supplement 1C). This value represents a wider range than that in (A). (D) Local accumulation similar to intercellular distribution of mV-Wnt8 and -Frzb. *a*_n,max_ is given randomly for each *n* by an absolute value of the normal distribution. (E) Distant scaffolds from the source region. *a*_n,max_ is given to depend on space: 10.0 for 50 ≤ *x* ≤ 60, otherwise, *a*_n,max_ is 0.0. This situation is similar to tethered-anti-HA Ab (Figure 2B). (F) Ligand accumulation in front of the HS-absent region. *a*_n,max_ is given depending on space: 10.0 for 0 ≤ *x* ≤ *L*/10 and 0.0 for *L*/10 < *x*. The values of *v* in (B) (*v*_B_) are also shown in grey, for comparison. Note that ligand accumulation occurs in front of the HS-absent region (*L*/10 < *x*.).

Under a wide range of appropriate parameter values, distributions of *u* and *v* converged to steady states within a few minutes (Figure 5B). The free component, *u*, quickly decreases, displaying a shallow continuous distribution pattern due to diffusion. In contrast, the bound component, *v*, shows a discrete distribution following *a*(*x*), and the level of *v* is much higher than *u* at any position, reflecting our conclusion that the majority of Wnt or sFRPs molecules in the extracellular space are bound. Given that activation of Wnt signaling requires internalization of the ligands (Kikuchi et al., 2009; Yamamoto et al., 2006) of the bound component, corresponding to *cv* in equation 3, the distribution of the bound component in this model could be equivalent to the “actual” gradient of Wnt signaling, even though it is not diffusing. We demonstrated that consistently some portion of Wnt8 ligands accumulated on *N*-sulfo-rich HS clusters initiate canonical Wnt signaling by forming the signaling complex “signalosome” (Mii et al., 2017).

We have shown that *N*-sulfo-rich HS, but not *N*-acetyl-rich clusters, are frequently endocytosed (Mii et al., 2017). In this model, different internalization rates of *N*-acetyl-rich and *N*-sulfo-rich HS clusters can be reflected by varying the internalization rate of the docking sites, *c*. A smaller value of *c* results in a long-range distribution (compare Figures 5B and 5C), explaining why Frzb shows a long-range distribution by binding to *N*-acetyl-rich HS clusters (Mii et al., 2017). We can evaluate the distribution by the decay length, *λ*. *λ* represents a distribution range when the steady state gradient is written as

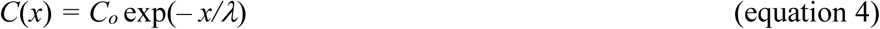

(Kicheva et al., 2012; Kicheva et al., 2007). We calculated *λ* by curve fitting the peak values of *v* to equation 4. The value of *λ* with *c* = 0.1 or 0.01 is 6.35 or 10.8 μm, respectively, showing that internalization rates of HS clusters can affect distribution ranges, as observed between Wnt and sFRPs (Mii and Taira, 2009). In addition, we examined the contribution of dissociation from the bound to the diffusing state, suggested by our cell-boundary FDAP (Figure 4). Without dissociation, a shorter-range distribution of the bound component was obtained (*λ* = 4.50 μm, Figure 5-Figure supplement 1) than with dissociation (*λ* = 6.35 μm, same data as in Figure 5B). Furthermore, in *Xenopus* embryos, Wnt8 in the intercellular space exhibited local accumulations (Figure 1B). In our model, when the binding rate *a*_n,max_ (equation 7 in the Materials and Methods) fluctuates randomly (i.e. the amount of HS at position *x*), the bound component of ligands, also fluctuates (Figure 5D), reproducing the local accumulation of Wnt8 and Frzb in *Xenopus* embryos. Even under these conditions, the free component shows a continuously decreasing gradient (Figure 5D, inset), which probably corresponds to the FCS-measured, diffusing component of the FGF8 gradient (Yu et al., 2009). Thus, our mathematical model can generalize protein distributions in the extracellular space.

## DISCUSSION

As one of major secreted signaling molecules, mechanisms of Wnt dispersal are crucial when we consider embryonic patterning and various other systems involving Wnt signaling (Routledge and Scholpp, 2019). Among many Wnt proteins, Wg distribution in the *Drosophila* wing disc has long been investigated as a morphogen gradient (Strigini and Cohen, 2000; Zecca et al., 1996). Various genetic studies show that the extracellular distribution of Wg largely depends on HSPGs, such as Dally and Dally-like glypicans (Baeg et al., 2004; Franch-Marro et al., 2005; Han et al., 2005). Furthermore, FRAP-based analysis suggests that the effective diffusion coefficient of Wg is much slower (0.05 µm^2^/s) than free diffusion (> 10 µm^2^/s) (Kicheva et al., 2007). However, such dynamics of secreted signaling proteins still remain a matter of debate (Rogers and Schier, 2011). On the other hand, recently we found that HS chains on the cell surface are organized in clusters with varying degrees of *N*-sulfo modification in *Xenopus* embryos and HeLa cells. Furthermore, we demonstrated that endogenous Wnt8 protein visualized by immunostaining shows a punctate distribution, specifically associated with *N*-sulfo-rich HS clusters (Mii et al., 2017). Similar punctate distributions have also been observed with Wg in *Drosophila* (Strigini and Cohen, 2000; van den Heuvel et al., 1989), but the significance of this distribution has not yet been explained. Therefore, to gain insight into the mechanism of Wnt distribution, we examined Wnt8 protein dynamics.

Based on quantitative live-imaging techniques, we propose that most Wnt8 molecules distributed among cells are mostly cell-surface-bound, while several percent of them are diffusing. Of note, we derived similar ratios of cell-surface-bound and diffusing molecules using two completely different methods (Figures 3G and 4E). In recent studies, small populations of mobile/diffusing molecules have commonly been observed with Wnt/EGL-20 in *C. elegans* (Pani and Goldstein, 2018) and Dpp in *Drosophila* wing disc (Zhou et al., 2012). Importantly, it has been suggested that these populations disperse over long distances, similar to our observation of mV-Wnt8 trapped using morphotrap (Figure 2C, D), generalizing the existence of long-dispersing populations in various model systems. Thus, to understand the mechanism of Wnt dispersal, it seemed important to determine how the bound/diffusing ratio is regulated in tissues.

Based on our previous observations, it is plausible that cell-surface-bound Wnt8 is mostly associated with HS clusters (Mii et al., 2017). The function of HSPGs in Wnt dispersal has been examined by genetic studies of *Drosophila.* These studies show that HSPGs are required for accumulation and transfer of Wnt ligands. Based on these results, it has been proposed that Wnt disperses by restricted diffusion, in which HSPGs transfer Wnt ligands in a bucket brigade manner (Yan and Lin, 2009). In contrast, results of this study seem inconsistent with this model. First, our FDAP assay showed that most photoconverted mK-Wnt8 does not diffuse laterally, even when other puncta of Wnt8 exist near the site of photoconversion (Figure 4-Figure supplement 2C, left panel, Video 1). Second, our morphotrap assay is incompatible with restricted diffusion model. In our morphotrap assay, mV-Wnt8 can move as much as 10-15 cell diameters (at least 200 µm) within 24 hours. However, if we consider restricted diffusion with a slow effective diffusion coefficient (0.05 µm^2^/s), 4 x 10^5^ sec (over 4 days) is required until the mean squared displacement (MSD, 2*Dt* in a one-dimensional space) of the molecule reaches a value corresponding to 10 cell diameters (200^2^ µm^2^). In contrast, if we consider free diffusion (*D* = 20 µm^2^/s), only 10^3^ sec is required for the same MSD. Therefore, freely diffusing Wnt ligands observed with FCS (fast component in Figure 3F) seem necessary to explain the long-range dispersal observed in our morphotrap assay. Thus, our results are inconsistent with the restricted diffusion model. Therefore, we propose a model in which once a Wnt ligand is released from an HS cluster (Figure 4), it diffuses freely until it binds to another HS cluster. Given that HS clusters are discretely distributed on cell surfaces, it is highly probable that locally accumulated Wnt ligands on HS clusters move to other clusters by changing from a bound state to a diffusing one. Therefore, we propose that the discrete distribution of HSPGs contributes to efficient, long-range dispersal of Wnt proteins in tissues.

Notably, in our mathematical model, distributions of both free and bound components converged to steady states within a few minutes, showing rather fast dynamics in the context of embryonic patterning. This characteristic could solve perceived weaknesses of diffusion-base models (Muller et al., 2013), especially dilemmas related to the speed and stability of gradient formation. From this point of view, the combination of abundant cell-surface-bound and minimal diffusing populations would be beneficial for signaling stability and speed of pattern formation, respectively.

Our cell-boundary FDAP suggests that cell-surface-bound and diffusing populations are probably exchangeable. Although this result can be explained by dissociation of molecules from the bound state as described above, it also seems possible that endocytosis reduces the number of photoconverted molecules (Figure 4C). However, we consider this less likely. Endocytosis of Wnt8 is possibly mediated by caveolin (Mii et al., 2017; Yamamoto et al., 2006), and we have already shown that internalized Wnt8 was detected as puncta in the cell (Mii et al., 2017). However, in FDAP analyses in this study, we excluded this possibility when we detected internalization of Wnt puncta in cells before curve-fitting analysis. In our mathematical model, when dissociation from the cell-surface does not occur (*b* = 0 in equation 2 and 3, Figure 5A), the range of the gradient (decay length, *λ*) was shortened from 6.35 to 4.50 μm (Figure 5-Figure supplement 1). Thus, at least in cases we analyzed, dissociation from the bound state seems to contribute to the long-range distribution and rapid formation of the gradient.

FRAP/FDAP and FCS are widely used to analyze dynamic states of morphogens (Kicheva et al., 2012; Rogers and Schier, 2011). Both methods can measure diffusion coefficients (*D*) of fluorescent protein-tagged morphogens. However, reported values for *D* by FRAP/FDAP and FCS often differ significantly (Kicheva et al., 2012; Kicheva et al., 2007; Yu et al., 2009). Moreover, comparisons of *D* values from FRAP/FDAP and FCS may not be relevant, because FRAP/FDAP usually examines a wider area and employs a longer analysis time than FCS (Rogers and Schier, 2011). To develop a comprehensive understanding of these methods and their results, we introduced a novel analysis for FCS and FDAP. Although FCS has single-molecule sensitivity, it cannot measure immobile molecules (Hess et al., 2002; Zhou et al., 2012). To complement this method, we used photon-counting imaging with a confocal microscope with a long pixel dwell time, which enabled us to detect even freely diffusing proteins (Supplementary Note). Using this complementary information, we estimated that the diffusing population comprises only several percent (Figure 3G). In order to design FDAP measurements that are comparable with FCS, we performed intercellular FDAP (Figure 4). Because we did not observe lateral accumulation of photoconverted mKikGR-Wnt8 (Figure 4—Figure supplement2C), it is better to use the binding/dissociation model (equation 1) rather than diffusion-based models, which consider a single diffusing component with an effective/apparent diffusion coefficient of *D_a_*. However, in order to compare our data with those previously reported (Kicheva et al., 2012), we also performed curve-fitting with an effective/apparent diffusion model (Figure 4-Figure supplement3, equation 5). As a result, the apparent diffusion coefficient *D_a_* (μm^2^/s) was calculated as 0.042 and 0.059 for mKikGR-Wnt8 and mKikGR-Frzb, respectively. These values are very close to a previously reported FRAP value for Wg (0.05 μm^2^/s) (Kicheva et al., 2007). Thus, such small values of *D_a_* relative to free diffusion could be interpreted as a result of interaction with cell surfaces, regardless of whether the protein of interest actually shows lateral diffusion in bucket brigade fashion. Nevertheless, as discussed above, these small *D_a_* values cannot explain the results of the morphotrap assay. We propose that the large difference in diffusion coefficients between FRAP/FDAP and FCS can be reconciled, if we consider the coexistence of a large immobile population and a tiny diffusing population (Figure 4F). Thus, FRAP/FDAP and FCS are likely to observe different populations of a secreted protein, even when analyzed on comparable spatio-temporal scales.

Based on our data, we propose a mathematical model consisting of free and bound components of Wnt (Figure 5A). This model can be widely applied to secreted proteins that bind to cell surfaces, including sFRPs and other peptide growth factors. Like tethered-anti-HA Ab (Figure 2B), atypical distributions of FGF (Shimokawa et al., 2011) and Nodal (Marjoram and Wright, 2011), in which ligands accumulate in locations distant from their sources, have been reported, although a theoretical explanation of these atypical distributions has proven elusive. In our model, the atypical distributions can be reproduced if specific scaffolds for ligands (ligand binding proteins) are anchored on the surfaces of cells (Figure 5E). Furthermore, our model explains the puzzling localization of ligands in tissues. In mosaic analyses of the wing discs of *Drosophila* mutants, Hh and Dpp ligands accumulate at the edges of clones defective in HS synthesis (Takei et al., 2004; Yan and Lin, 2009). Distributional patterns of these ligands are explained by our model, which accounts for accumulations of ligand in regions lacking HS (Figure 5F). Thus, our model provides comprehensive understanding of the extracellular behavior of secreted proteins.

## MATERIALS AND METHODS

All experiments using *Xenopus laevis* were approved by The Institutional Animal Care and Use Committee, National Institutes of Natural Sciences. or The Office for Life Science Research Ethics and Safety, University of Tokyo.

### *Xenopus* embryo manipulation and microinjection

Unfertilized eggs of *Xenopus laevis* were obtained by injection of gonadotropin (ASKA Pharmaceutical). These eggs were artificially fertilized with testis homogenates and dejellied by 4% L-cysteine solution (pH adjusted to 7.8 by addition of NaOH). Embryos were incubated in 1/10x Steinberg’s solution at 14-17 °C and were staged according to Nieuwkoop and Faber (Nieuwkoop and Faber, 1967). Synthesized mRNAs were microinjected into early (2-16 cells) embryos. The amounts of injected mRNAs are described in the figure legends.

### Fluorescent image acquisition

Image acquisition was performed using confocal microscopes (TSC SP8 system with HC PL APO ×10/NA0.40 dry objective or HC PL APO2 ×40/NA1.10 W CORR water immersion objective, Leica or LSM710 system with C-Apochromat 40x/1.2 W Corr M27 water immersion objective, Zeiss). Photon counting images were acquired with a HyD detector (Leica). Detailed conditions for the imaging are available upon request. mV was constructed by introducing an A206K mutation to prevent protein aggregation (Zacharias et al., 2002). mV or mK-tagged constructs were shown to be biologically active (Mii et al., 2017) (Figure 4-Figure supplement 1). For FDAP and FCS measurements, gastrula embryos were embedded on 35 mm glass-based dishes (Iwaki) with 1.5% LMP agarose (#16520-050; invitrogen) gel, which was made of 1/10x Steinberg’ solution. For other live-imaging, embryos were mounted in house-made silicone chamber with holes of 1.8 mm-diameter. Fluorescent intensity was measured using Fiji, Image J (NIH) or Zen2009 (Zeiss).

### cDNA cloning of IgG from cultured hybridomas

Cultured hybridomas were harvested by centrifuge and total RNAs were prepared using ISOGEN (Nippon Gene), according to the manufacturer’s protocol. First strand cDNA pools were synthesized using SuperScript II reverse transcriptase (Invitrogen) and random hexamer oligo DNA. These cDNA pools were used as templates for PCRs to isolate cDNAs for heavy chains and light chains of anti-HA and anti-Myc IgGs. See Table S1 for all primers used for PCR cloning. The full length cDNAs were cloned into the pCSf107mT vector (Mii & Taira, 2009).

Cultured hybridoma cells were harvested by centrifuge and total RNAs were prepared using ISOGEN (Nippon Gene), according to the manufacturer’s protocol. First strand cDNA pools were synthesized using SuperScript II reverse transcriptase (Invitrogen) and random hexamer oligo DNA. These cDNA pools were used as templates for PCRs to isolate cDNAs for heavy and light chains of anti-HA and anti-Myc IgGs. Procedures of PCR cloning were as follows. IgG cDNAs for 3’ regions of CDSs were obtained by PCR with degenerate primers (5’ γ-F and 5’ κ-F) and primers corresponding to constant regions of Ig genes (γ2b-const-R, γ1-const-R, 3’ κ-R) (Wang et al., 2000). To obtain the complete CDSs, 5’RACE was carried out to obtain the first codons of Ig genes, by using a modified protocol, in which inosines are introduced into the G-stretch of the HSPst-G10 anchor (personal communications from Dr. Min K. Park). cDNAs were synthesized with gene specific primers (HA-heavy-R1, Myc-heavy-R1, HA-light-R1, and Myc-light-R1), and tailed with poly-(C) by terminal deoxynucleotidyl transferase, and subsequently double strand cDNAs were synthesized with the HSPst-G10 anchor. The 5’ ends of cDNAs were amplified by PCR between the HSPst adaptor and gene-specific primers (HA-heavy-R2, Myc-heavy-R2, HA-light-R2 and Myc-light-R2) using the double strand cDNAs as templates. Full length CDSs were amplified using primers designed for both ends of the CDSs (HH-Bam-F, MH-Bam-F, HL-bam-F, and ML-Bgl-F for 5’ ends; 3’ γ2b-R, 3’ γ1-R and 3’ κ-R for the 3’ end) and the first cDNA pools. See Table S1 for all primers used for PCR cloning. The full length cDNAs were cloned into the pCSf107mT vector (Mii and Taira, 2009).

### FDAP measurements

For expression in the animal cap region of *Xenopus* embryos, four cell-stage *Xenopus laevis* embryos were microinjected with mRNAs for mK-Wnt8 and mK-Frzb (4.0 ng/embryo) at a ventral blastomere. Injected embryos were incubated at 14 °C until the gastrula stage (st. 10.25-11.5) for subsequent confocal analysis. FDAP measurements were performed using the LSM710 system (Zeiss) with a C-Apochromat ×40, NA1.2 water immersion objective. Time lapse image acquisition was carried out for 20 seconds each at 25 frames/s, and after 4 seconds (100 frames) from the start, intercellular mK-fusion proteins were photoconverted at a small rectangular region (1.66 × 2.49 μm) with 405 nm laser light irradiation. After the photoconversion, images were acquired for 16 seconds (400 frames). Red fluorescent intensities within the rectangular region where photoconversion was performed, were analysed by curve-fitting to equation 1 (Figure 4D) or equation 5 (Figure 4-Figure supplement 3A), using the Curve Fitting Toolbox of MATLAB (Mathworks).

### FCS measurements

FCS measurements were carried out using a ConfoCor2 system (Zeiss) with a C-Apochromat ×40, NA1.2 water immersion objective, according to a previous report (Pack et al., 2006). mRNAs for mV-Wnt8 and sec-mV were microinjected into 4-cell stage Xenopus embryos. Amount of mRNAs were both 250 pg/embryo. Injected embryos were measured at gastrula stage (st. 10.25-11.5). Places where measurements were performed were determined by bright field view, to confirm the places were extracellular.

### Plasmid construction

pCSf107mT (Mii and Taira, 2009) was used to make most plasmid constructs for mRNA synthesis. pCSf107SPw-mT and pCSf107SPf-mT were constructed, which have the original signal peptides of Wnt8 and Frzb, respectively. The coding sequence (CDS) for mVenus (mV) or mKikGR (mK) was inserted into the BamHI site of pCSf107SPw-mT or pCSf107SPf-mT to construct pCSf107SPw-mV-mT, pCSf107SPf-mV-mT, pCSf107SPw-mK-mT, and pCSf107SPf-mK-mT. Constructs for SP-mV, SP-mV-HB, and SP-mV-2HA were made with pCSf107SPf-mT. pCS2+HA-IgH-TM-2FT (the heavy chain for anti-HA IgG with the transmembrane domain of a membrane-bound form of IgG heavy chain) was made by inserting the full length CDS of heavy chain of anti-HA IgG without the stop codon (using the EcoRI and BglII sites) and a partial CDS fragment corresponding to the IgG transmembrane domain (using the BglII and XbaI sites) into the EcoRI/XbaI sites of pCS2+mcs-2FT-T.

### Luciferase reporter assays

Luciferase reporter assays were carried out as previously described (Mii and Taira, 2009). Multiple comparisons were carried out with pairwise Wilcoxon rank sum test (two-sided) in which significance levels (*p*-values) were adjusted by the Holm method using R.

### Mathematical modeling

Two ligand components were considered: free and bound ones. This model includes five dynamic processes: (i) the production of ligands, (ii) the diffusion of the free component in the intercellular space, (iii) the binding of ligands to dotted structures (“docking sites”) such as HS clusters on the surface of cells, (iv) the release of the bound component from the “docking sites,” and (v) the internalization of the bound component into cells. The model in one-dimensional space is written as:

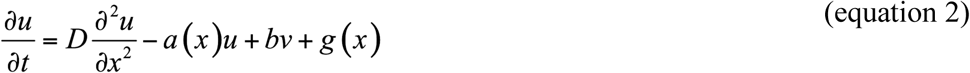

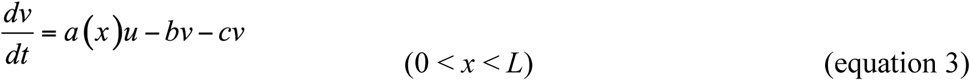

where *u* and *v* represent the amounts of the free and bound components of morphogen molecules, respectively. The symbols *a*(*x*), *b*, *c*, and *g*(*x*) represent binding, release, internalization, and production rates, respectively. *D* represents the diffusion coefficient of the free component in the extracellular space. The ligand is assumed to be produced in a limited region using the following function:

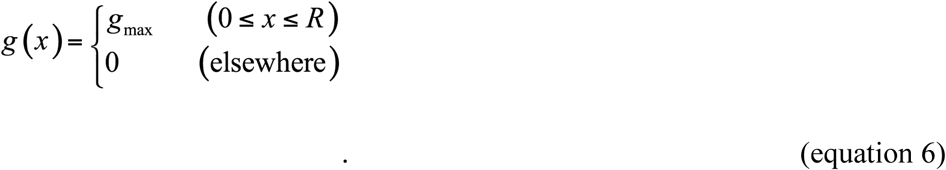

We assumed that the binding rate *a*(*x*) depends on the position *x*, following heterogeneous distribution of HS clusters on the cell surface. The following function was used for *a*(*x*):

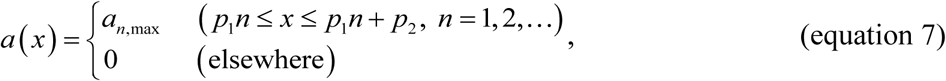

where *p_1_* and *p_2_* are the interval and width, respectively, of docking sites. We used the no-flux (Neumann) boundary condition at *x* = 0 and *L*. We calculated the model by numerical simulation. The initial distribution of *u* and *v* was set at 0 in the entire space.

In the one-dimensional space, distributions of free (*u*) and bound (*v*) components of secreted proteins were obtained by computer simulation, where the spatial length *L* = 100 (µm). Distributions of *u* (red) and *v* (blue) are presented at time *t* = 100 (sec), which almost reached steady states. We used the forward difference method with the spatial step Δ*x* = 0.1 and temporal step Δ*t* = 0.0001 in numerical calculations. In Figure 5B, parameter values are: *D* = 20.0, *a*_n,max_ = 10.0, *b* = 0.1, *c* = 0.1, *g*_max_ = 0.2, *R* = *L*/100, *p*_1_ = 2, and *p*_2_ = 0.2. In other panels, distributions in specific conditions are shown (see figure legends).

## ACKNOWLEDGEMENTS

We thank Dr. Min Kyun Park for hybridoma culture and 5’RACE for IgG cDNAs. We also thank Dr. A. Miyawaki for Venus cDNA; Dr. R. Moon for pCS2+Xwnt8; Dr. Shinya Matsuda for critical reading. This work was supported in part by following programs: KAKENHI (24870031, 18K14720), the NINS program for cross-disciplinary study (1311608, 01311801), JST-PRESTO (JPMJPR194B).

## Author contributions

Yusuke Mii, Conceptualization, Formal analysis, Investigation, Writing-original draft, Writing-review and editing; Kenichi Nakazato, Mathematical Modeling-design and writing codes; Chan-Gi Pack, FCS analyses, Writing-review and editing; Yasushi Sako, Supervision of FCS analyses; Atsushi Mochizuki, Mathematical Modeling-design and supervision; Shinji Takada, Supervision, Writing-review and editing; Masanori Taira, Supervision, Writing-review and editing.

## VIDEO LEGENDS

**Videos 1 and 2. Photoconversion of mKikGR-Wnt8 in a cell-boundary region of the *Xenopus* embryo.**

**Video 3. Photoconversion of mKikGR-Frzb in a cell-boundary region of the Xenopus embryo.**

Photoconversion of mKikGR fusion proteins was performed at a cell-boundary region in animal cap of *Xenopus* gastrula (st. 10.5-st.11.5). mKikGR-Wnt8 (Videos 1 and 2) or mKikGR-Frzb (Video 3) was photoconverted at the region indicated with the blue box after 100 flames scanned (about 4 seconds), and further 400 flames were scanned for measurement. The width of the region for photoconversion and intensity measurement was 20 pixels (1.66 μm). The play speed is x1.

## SUPPLEMENTARY TABLE

**Table 1.**
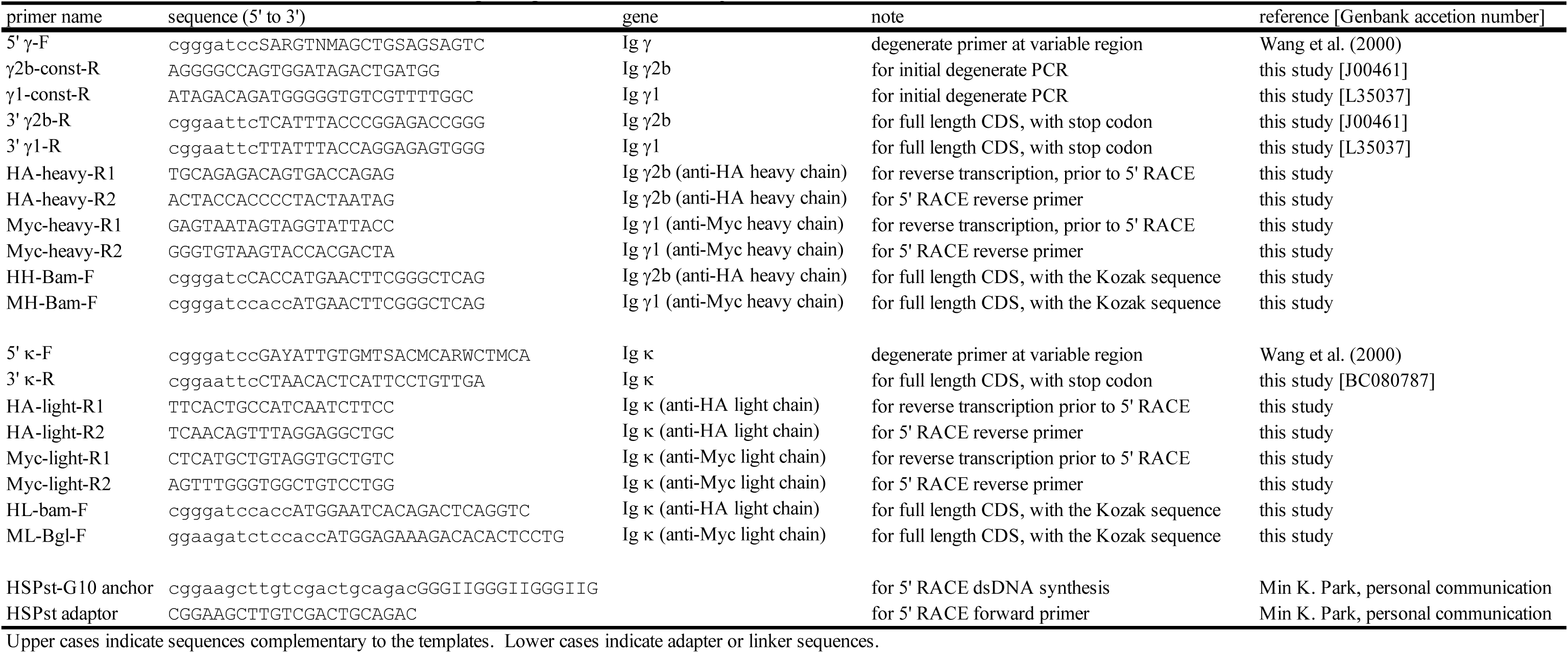
Primers used for molecular cloning of IgG cDNAs from hybridomas. See the section of “cDNA cloning of IgG from cultured hybridomas” in Metarials and Methods for detail.

## SUPPLEMENTARY NOTE

### Detecting freely diffusing molecules with confocal microscopy

When we consider detection of diffusing molecules with confocal microscopy, diffusing molecules may not be detected because of their large displacement. In our imaging conditions, pixel dwell time was 24.4 μs and observed maximum *D_fast_* in FCS was about 80 μm^2^/s (Figure S3E). During pixel dwell time *t*, mean square displacement (MSD) of molecules of diffusion coefficient *D* in three-dimensional space is 6*Dt* (Crank, 1975). Therefore, maximum MSD is estimated as

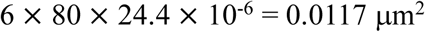

Accordingly, mean displacement of the molecules at the maximum is

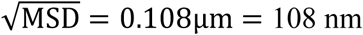

On the other hand, when the numerical aperture (*NA*) of an objective lens is given, the Reyligh diffraction limit in our condition is calculated as

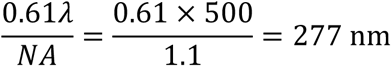

This value of the diffraction limit is larger than the 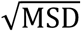 so that even the most rapidly diffusing molecules observed in FCS measurements are detectable using confocal microscopy.

---

**Figure 1-Figure supplement 1.**
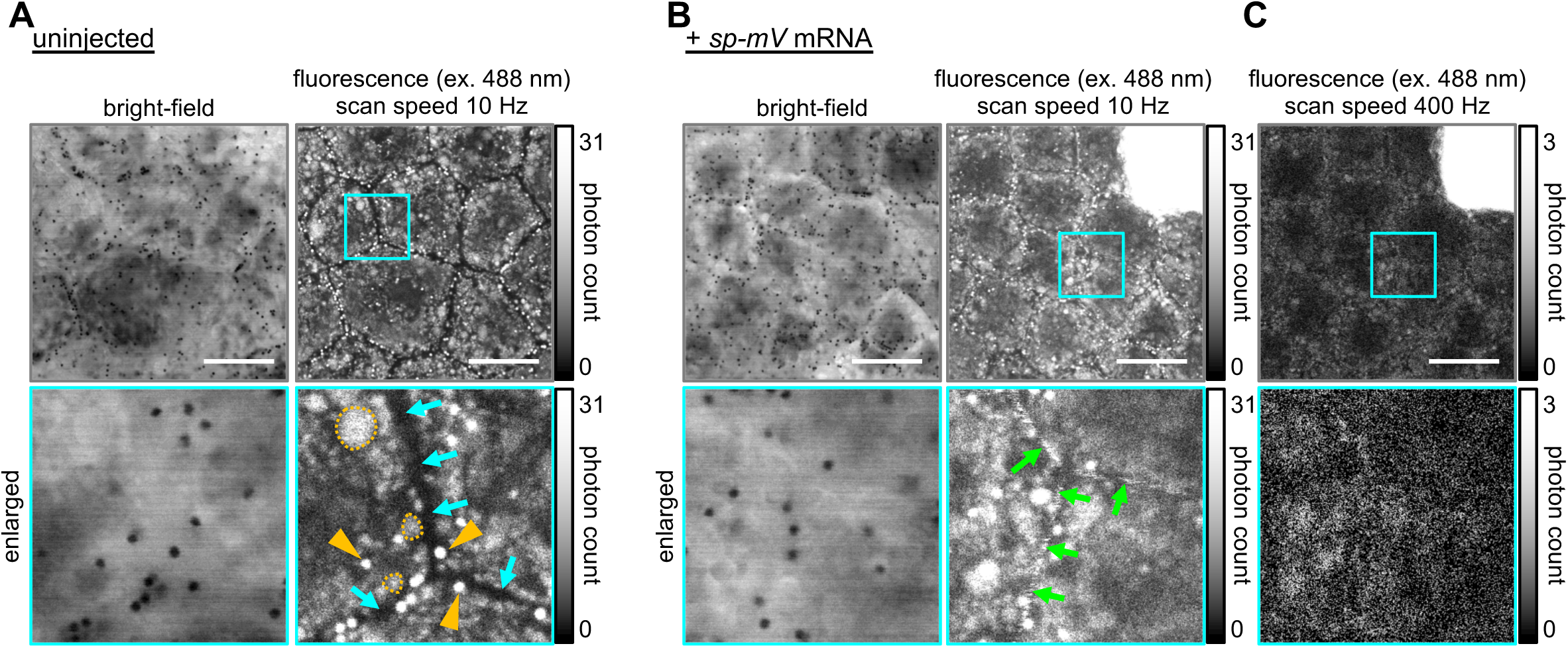
Imaging of secreted mVenus protein in the intercellular space. All images presented were acquired using live-imaging with photon counting detection. LUT used is as indicated. (A) Photon counting image of uninjected gastrula. In such a highly enhanced condition, autofluorescence from pigment granules (orange arrowheads) and yolk granules (orange dashed circles) can be observed. However, little fluorescence was observed in the intercellular region (cyan arrows). The original data was the same as in Figure 1A. (B, C) Photon counting image of *sp-mV* mRNA injected gastrula. (B) Data acquisition and processing was done in the same condition as A. Unlike uninjected embryos, fluorescence signal was detected in the intercellular region (green arrows), indicating existence of secreted mVenus (sec-mV) protein. The original data was the same as in Figure 1A. (C) A faster scan speed resulted in an image of poor quality. The same position as in B was scanned at 400 Hz (this is the default setting of the Leica SP8 confocal system). In this condition, fluorescence in the intercellular space was hard to recognize. Scale bars, 20 μm. Amounts of injected mRNAs (pg/embryo): *sp-mV*, 250.

**Figure 2-Figure supplement 1.**
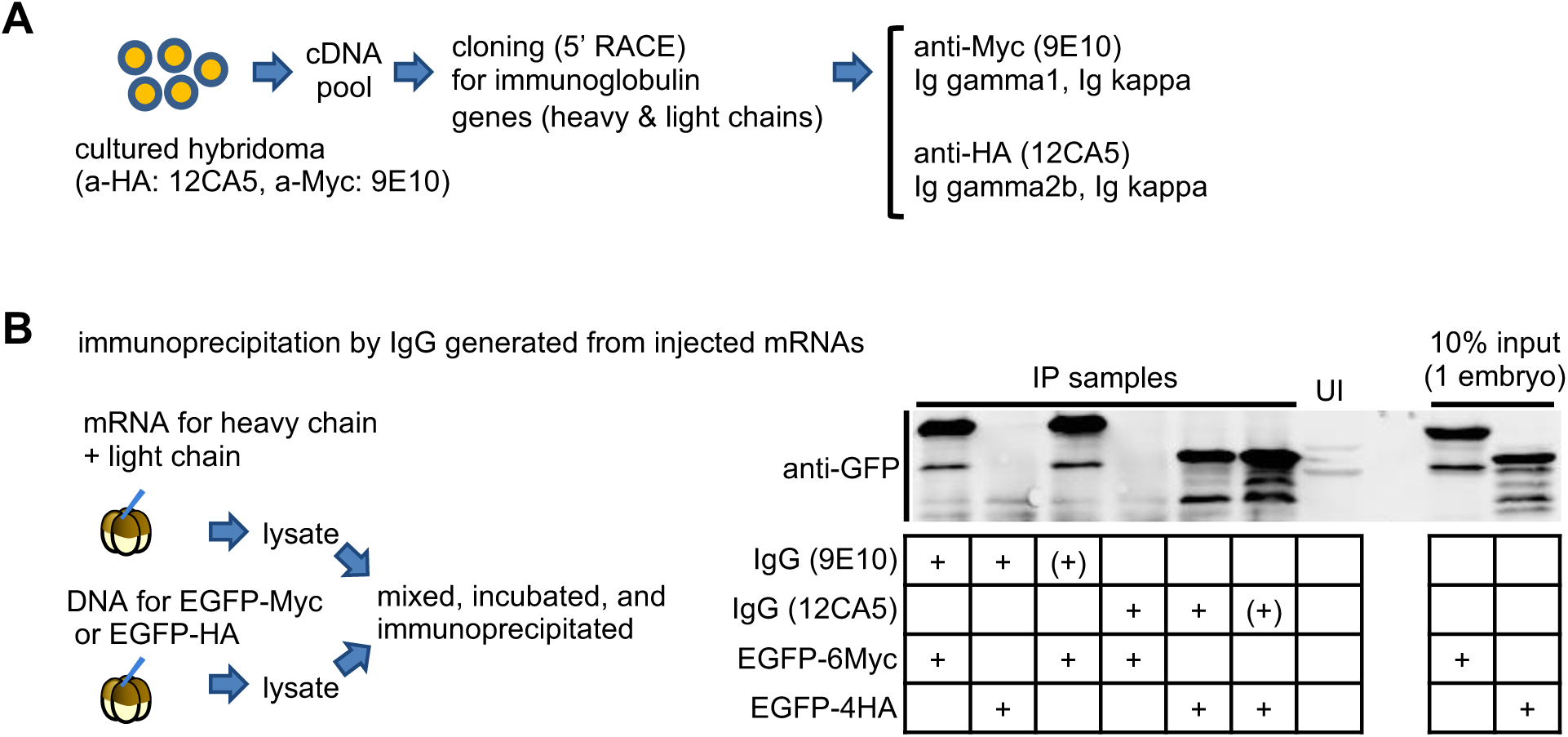
Generation of functional antibody protein by mRNA-injection into Xenopus embryo. (A) Summary of cDNA cloning for anti-HA and anti-Myc monoclonal antibodies. (B) Immunoprecipitation assays to examine the specificity of IgG (9E10, anti-Myc; 12CA5, anti-HA) generated from injected mRNAs. Combinations of lysates are indicated by +. (+) indicates the addition of the monoclonal antibody as a positive control. IgG generated in *Xenopus* embryos properly worked, similar to the monoclonal antibody. UI, lysate of uninjected embryos. EGFP-6Myc and EGFP-4HA were expressed by DNA injection (50 ng/embryo). Amounts of injected mRNAs (ng/embryo): heavy chains (B), 1.0; light chains (B), 0.61.

**Figure 3-Figure supplement 1.**
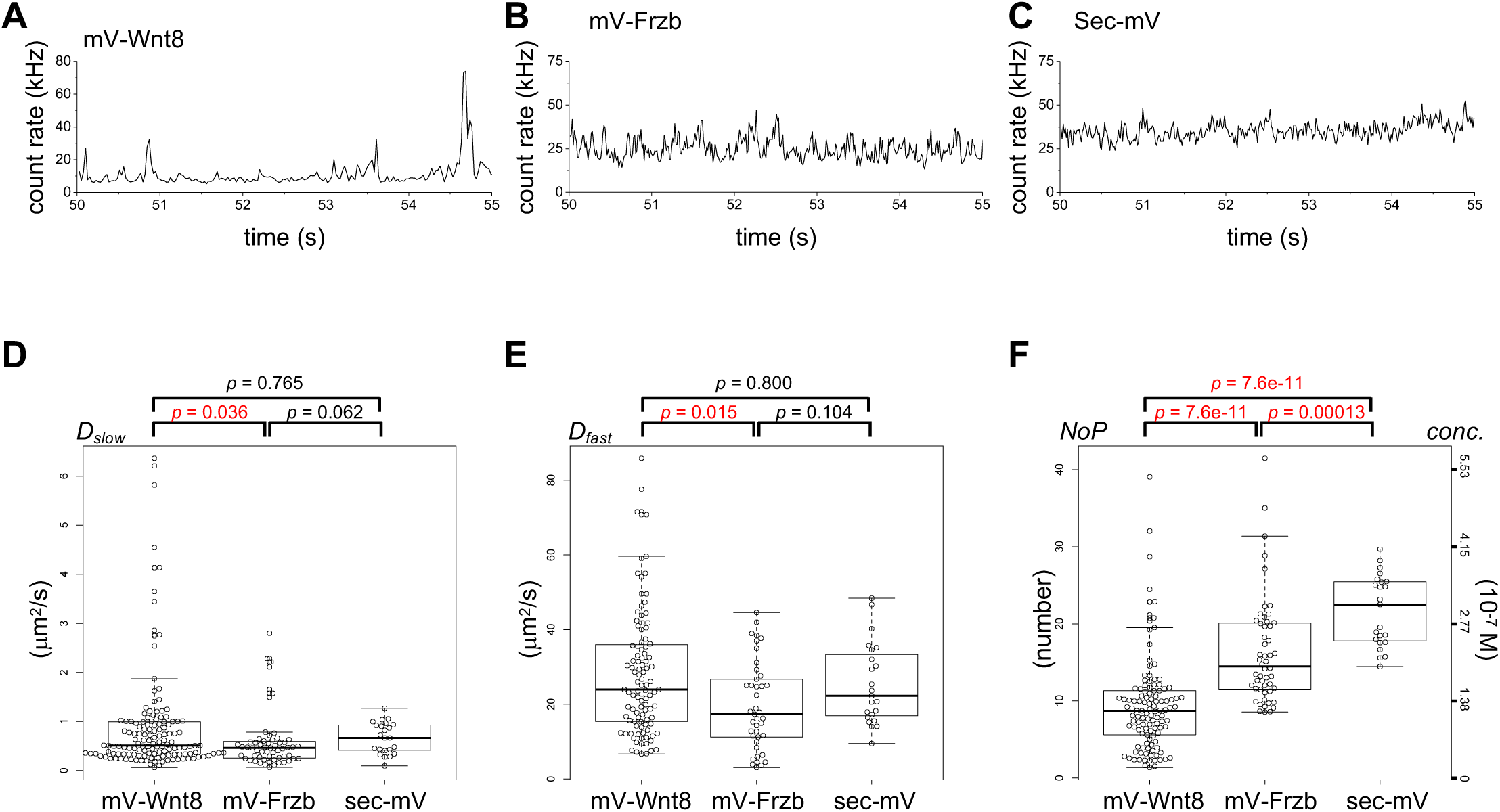
Supplementary data for FCS analysis. (A-C) Time course of fluorescent intensities for mV-Wnt8 (A), mV-Frzb (B) and sec-mV (C). Magnified views in a 5-second window as indicated by blue double headed arrows in Figure 3B-D. (D-F) Distribution of *D_slow_*, *D_fast_*, and number of particles (*NoP*) in individual measurements of FCS. Dot plots for individual data values and boxplots were drawn with R and its package “beeswarm”. Statistical significance (*p*) was calculated by Wilcoxon rank sum test (two-sided) and when it was significant, the *p* value is indicated in red.

**Figure 4-Figure supplement 1.**
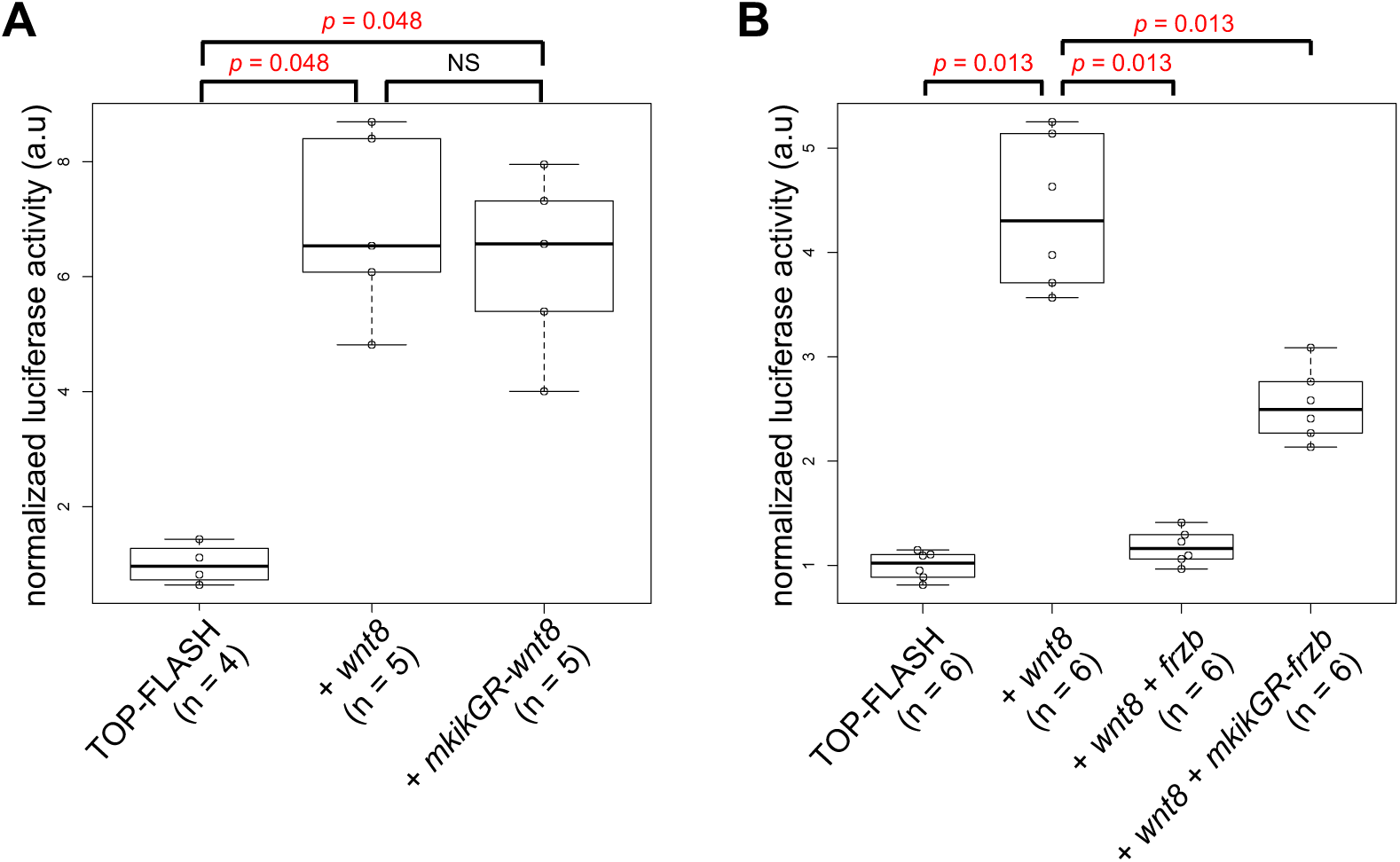
Biological activity of mKikGR-Wnt8 and -Frzb. TOP-FLASH reporter assay. The reporter DNA was injected into the animal pole region of a ventral blastomere of 4- or 8-cell stage *Xenopus* embryos with or without mRNAs as indicated, and the injected embryos were harvested at st. 11.5. Dot plots for individual data values (normalized with the mean value of TOP-FLASH only) and boxplots were drawn with R and its package “beeswarm”. Number of a pool of three embryos for each sample are as indicated (n). Statistical significance (*p*) was calculated by Wilcoxon rank sum test (two-sided) and when it was significant, the *p* value is indicated in red. (A) *mkikGR-wnt8* showed a significantly higher activity than that of TOP-FLASH only, indicating the activation of Wnt signaling. (B) *mkikGR-frzb* showed a significantly lower activity than that of TOP-FLASH + wnt8, indicating the inhibition of Wnt signaling. Amounts of injected DNA/mRNAs (pg/embryo): TOP-FLASH DNA, 150; *wnt8* mRNA, 20; *mkikGR-wnt8* mRNA, 31 (equimoler to *wnt8*); *frzb* mRNA, 80; *mkikGR-frzb* mRNA, 127 (equimolar to *frzb*).

**Figure 4-Figure supplement 2.**
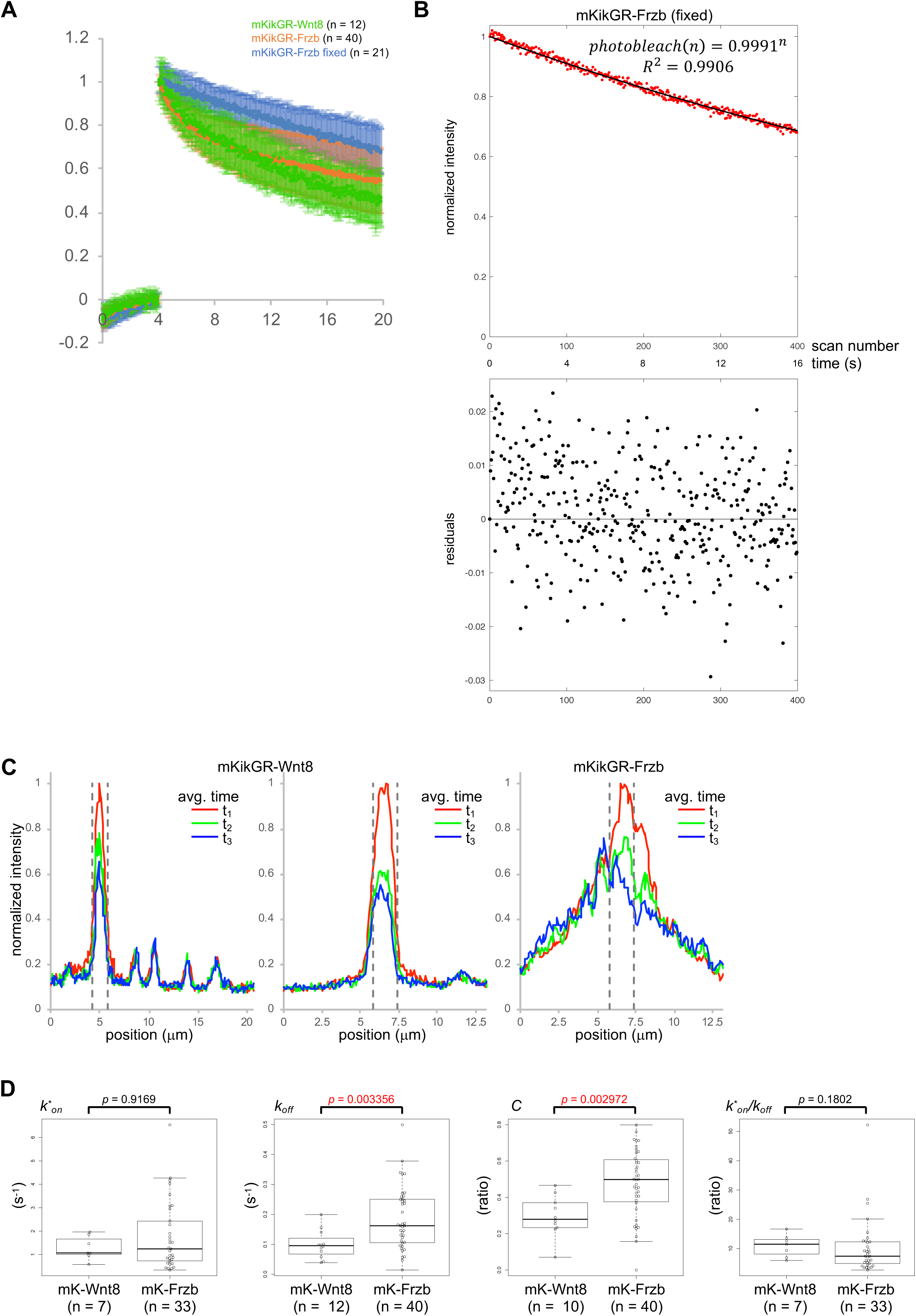
Fluorescence decay after photoconversion (FDAP) assay in the extracellular space of Xenopus embryos. (A) Time course of red (photoconverted state) fluorescent intensity within the photoconverted region. Means +/– standard deviations (s.d.) were presented. (B) FDAP of mKikGR-Frzb (fixed) fits with the photobleaching model caused by iterations of scanning. (C) Spatial intensity profiles of FDAP. Representative data were shown. Regions of photoconversion were indicated with vertical dashed lines. 1st panel corresponds to Video 1, showing no lateral accumulation even when punctate distributions of unconverted mK-Wnt8 proteins are present in the vicinity. 2nd and 3rd panels correspond to Videos 2 and 3, respectively. To reduce fluctuations of intensities, means of intensity data from ten flames were plotted for each time point (t_1_-t_3_). Averaged time after photoconversion (s): t_1_, 0.2178; t_2_, 7.9606; t_3_,15.6826. (D) Distribution of coefficients in individual measurements of FDAP obtained from fitting with the dissociation model. Dot plots for individual data values and boxplots were drawn with R and its package “beeswarm”. Statistical significance (*p*) was calculated by Wilcoxon rank sum test (two-sided) and when it was significant, the *p* value is indicated in red. Note that data with a large confidential interval (95% confidential interval > twice of the calculated value) were excluded due to their low reliability. As a result, numbers of measurement (n) was not always matched with those in Figure 4C.

**Figure 4-Figure supplement 3.**
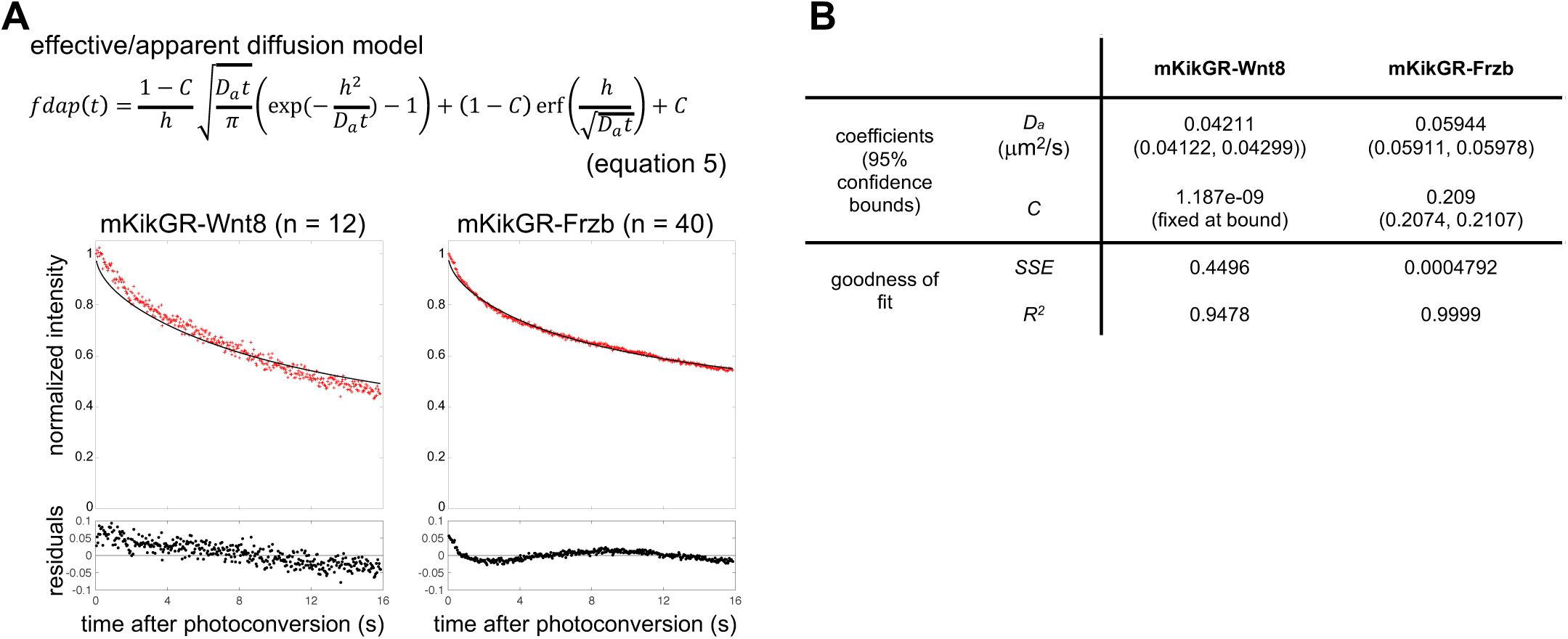
Fluorescent decay curve fitted with the effective diffusion model. (A) Fluorescent decay curve fitted with the effective diffusion model. The half-width of the photoconverted region is defined as *h* (= 0.83 μm) in equation 5. Mean of the normalized intensities for each time point was plotted with a red cross. Fitting curves are shown as black lines. The residuals were within 5% (0.05) in all cases. (B) Coefficients and evaluations of goodness of fit with the effective diffusion model. *D_a_*, apparent diffusion coefficient; *C*, ratio of immobile population; *SSE*, sum of squared errors; *R*^2^, coefficient of determination.

**Figure 5-Figure supplement 1.**
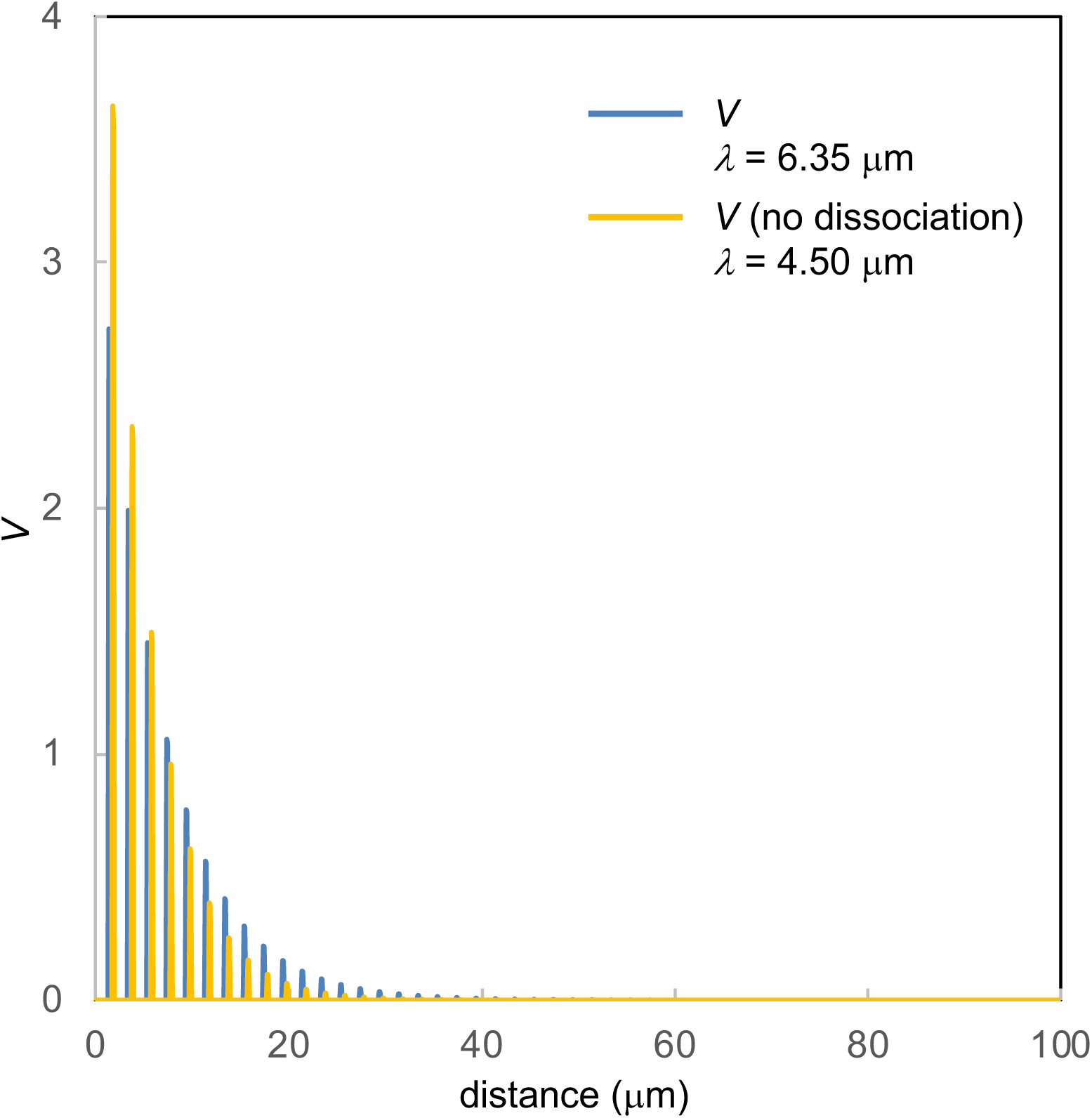
Contribution of dissociation from the HS-bound state to diffusing state. Distributions of the bound populations (*v*) are plotted. Blue line indicates results with dissociation from the bound state (the same data as Figure 5B). Yellow line indicates result without dissociation from the bound state (*b* = 0 in equations 2 and 3). Note that the position of the blue line is slightly moved to the left, to avoid overlapping with the yellow line. The decay length *λ* is as indicated.

## REFERENCES

Alexandre, C., Baena-Lopez, A., and Vincent, J. P. (2014). Patterning and growth control by membrane-tethered Wingless. Nature 505, 180–185.

Baeg, G. H., Selva, E. M., Goodman, R. M., Dasgupta, R., and Perrimon, N. (2004). The Wingless morphogen gradient is established by the cooperative action of Frizzled and Heparan Sulfate Proteoglycan receptors. Dev. Biol. 276, 89–100.

Chaudhary, V., Hingole, S., Frei, J., Port, F., Strutt, D., and Boutros, M. (2019). Robust Wnt signaling is maintained by a Wg protein gradient and Fz2 receptor activity in the developing Drosophila wing. Development 146.

Clevers, H., Loh, K. M., and Nusse, R. (2014). Stem cell signaling. An integral program for tissue renewal and regeneration: Wnt signaling and stem cell control. Science 346, 1248012.

Clevers, H., and Nusse, R. (2012). Wnt/beta-catenin signaling and disease. Cell 149, 1192–1205.

Crank, J. (1975). The mathematics of diffusion. 2nd ed., 2nd edn (Oxford [England]:, Oxford University Press).

Farin, H. F., Jordens, I., Mosa, M. H., Basak, O., Korving, J., Tauriello, D. V., de Punder, K., Angers, S., Peters, P. J., Maurice, M. M., and Clevers, H. (2016). Visualization of a short-range Wnt gradient in the intestinal stem-cell niche. Nature 530, 340–343.

Franch-Marro, X., Marchand, O., Piddini, E., Ricardo, S., Alexandre, C., and Vincent, J. P. (2005). Glypicans shunt the Wingless signal between local signalling and further transport. Development 132, 659–666.

Habuchi, S., Tsutsui, H., Kochaniak, A. B., Miyawaki, A., and van Oijen, A. M. (2008). mKikGR, a monomeric photoswitchable fluorescent protein. PLoS One 3, e3944.

Han, C., Yan, D., Belenkaya, T. Y., and Lin, X. (2005). Drosophila glypicans Dally and Dally-like shape the extracellular Wingless morphogen gradient in the wing disc. Development 132, 667–679.

Harmansa, S., Hamaratoglu, F., Affolter, M., and Caussinus, E. (2015). Dpp spreading is required for medial but not for lateral wing disc growth. Nature 527, 317–322.

Hess, S. T., Huang, S., Heikal, A. A., and Webb, W. W. (2002). Biological and chemical applications of fluorescence correlation spectroscopy: a review. Biochemistry 41, 697–705.

Kerszberg, M., and Wolpert, L. (1998). Mechanisms for positional signalling by morphogen transport: a theoretical study. J. Theor. Biol. 191, 103–114.

Kicheva, A., Bollenbach, T., Wartlick, O., Julicher, F., and Gonzalez-Gaitan, M. (2012). Investigating the principles of morphogen gradient formation: from tissues to cells. Curr. Opin. Genet. Dev. 22, 527–532.

Kicheva, A., Pantazis, P., Bollenbach, T., Kalaidzidis, Y., Bittig, T., Julicher, F., and Gonzalez-Gaitan, M. (2007). Kinetics of morphogen gradient formation. Science 315, 521–525.

Kiecker, C., and Niehrs, C. (2001). A morphogen gradient of Wnt/beta-catenin signalling regulates anteroposterior neural patterning in Xenopus. Development 128, 4189–4201.

Kikuchi, A., Yamamoto, H., and Sato, A. (2009). Selective activation mechanisms of Wnt signaling pathways. Trends Cell Biol. 19, 119–129.

Lin, X. (2004). Functions of heparan sulfate proteoglycans in cell signaling during development. Development 131, 6009–6021.

Loh, K. M., van Amerongen, R., and Nusse, R. (2016). Generating Cellular Diversity and Spatial Form: Wnt Signaling and the Evolution of Multicellular Animals. Dev. Cell 38, 643–655.

MacDonald, B. T., Tamai, K., and He, X. (2009). Wnt/beta-catenin signaling: components, mechanisms, and diseases. Dev. Cell 17, 9–26.

Marjoram, L., and Wright, C. (2011). Rapid differential transport of Nodal and Lefty on sulfated proteoglycan-rich extracellular matrix regulates left-right asymmetry in Xenopus. Development 138, 475–485.

Matsuda, T., Miyawaki, A., and Nagai, T. (2008). Direct measurement of protein dynamics inside cells using a rationally designed photoconvertible protein. Nat. Methods 5, 339–345.

Mii, Y., and Taira, M. (2009). Secreted Frizzled-related proteins enhance the diffusion of Wnt ligands and expand their signalling range. Development 136, 4083–4088.

Mii, Y., Yamamoto, T., Takada, R., Mizumoto, S., Matsuyama, M., Yamada, S., Takada, S., and Taira, M. (2017). Roles of two types of heparan sulfate clusters in Wnt distribution and signaling in Xenopus. Nat. Commun. 8, 1973.

Muller, P., Rogers, K. W., Jordan, B. M., Lee, J. S., Robson, D., Ramanathan, S., and Schier, A. F. (2012). Differential diffusivity of Nodal and Lefty underlies a reaction-diffusion patterning system. Science 336, 721–724.

Muller, P., Rogers, K. W., Yu, S. R., Brand, M., and Schier, A. F. (2013). Morphogen transport. Development 140, 1621–1638.

Nieuwkoop, P. D., and Faber, J. (1967). Normal Table of Xenopus laevis (Daudin) (Amsterdam, North Holland).

Nusse, R., and Clevers, H. (2017). Wnt/beta-Catenin Signaling, Disease, and Emerging Therapeutic Modalities. Cell 169, 985–999.

Pack, C., Saito, K., Tamura, M., and Kinjo, M. (2006). Microenvironment and Effect of Energy Depletion in the Nucleus Analyzed by Mobility of Multiple Oligomeric EGFPs. Biophys. J. 91, 3921–3936.

Pani, A. M., and Goldstein, B. (2018). Direct visualization of a native Wnt in vivo reveals that a long-range Wnt gradient forms by extracellular dispersal. eLife 7.

Rogers, K. W., and Schier, A. F. (2011). Morphogen gradients: from generation to inter-pretation. Annu. Rev. Cell Dev. Biol. 27, 377–407.

Routledge, D., and Scholpp, S. (2019). Mechanisms of intercellular Wnt transport. Development 146.

Shimokawa, K., Kimura-Yoshida, C., Nagai, N., Mukai, K., Matsubara, K., Watanabe, H., Matsuda, Y., Mochida, K., and Matsuo, I. (2011). Cell surface heparan sulfate chains regulate local reception of FGF signaling in the mouse embryo. Dev. Cell 21, 257–272.

Smith, J. C. (2009). Forming and interpreting gradients in the early Xenopus embryo. Cold Spring Harb. Perspect. Biol. 1, a002477.

Sprague, B. L., and McNally, J. G. (2005). FRAP analysis of binding: proper and fitting. Trends Cell Biol. 15, 84–91.

Sprague, B. L., Pego, R. L., Stavreva, D. A., and McNally, J. G. (2004). Analysis of binding reactions by fluorescence recovery after photobleaching. Biophys. J. 86, 3473–3495.

Strigini, M., and Cohen, S. M. (2000). Wingless gradient formation in the Drosophila wing. Curr. Biol. 10, 293–300.

Tabata, T., and Takei, Y. (2004). Morphogens, their identification and regulation. Development 131, 703–712.

Takada, R., Mii, Y., Krayukhina, E., Maruyama, Y., Mio, K., Sasaki, Y., Shinkawa, T., Pack, C. G., Sako, Y., Sato, C., et al. (2018). Assembly of protein complexes restricts diffusion of Wnt3a proteins. Commun. Biol. 1, 165.

Takei, Y., Ozawa, Y., Sato, M., Watanabe, A., and Tabata, T. (2004). Three Drosophila EXT genes shape morphogen gradients through synthesis of heparan sulfate proteoglycans. Development 131, 73–82.

van den Heuvel, M., Nusse, R., Johnston, P., and Lawrence, P. A. (1989). Distribution of the wingless gene product in Drosophila embryos: a protein involved in cell-cell communication. Cell 59, 739–749.

Verrecchio, A., Germann, M. W., Schick, B. P., Kung, B., Twardowski, T., and San Antonio, J. D. (2000). Design of peptides with high affinities for heparin and endothelial cell proteoglycans. J. Biol. Chem. 275, 7701–7707.

Wang, Z., Raifu, M., Howard, M., Smith, L., Hansen, D., Goldsby, R., and Ratner, D. (2000). Universal PCR amplification of mouse immunoglobulin gene variable regions: the design of degenerate primers and an assessment of the effect of DNA polymerase 3’ to 5’ exonuclease activity. J. Immunol. Methods 233, 167–177.

Yamamoto, H., Komekado, H., and Kikuchi, A. (2006). Caveolin is necessary for Wnt-3a-dependent internalization of LRP6 and accumulation of beta-catenin. Dev. Cell 11, 213–223.

Yan, D., and Lin, X. (2009). Shaping morphogen gradients by proteoglycans. Cold Spring Harb. Perspect. Biol. 1, a002493.

Yu, S. R., Burkhardt, M., Nowak, M., Ries, J., Petrasek, Z., Scholpp, S., Schwille, P., and Brand, M. (2009). Fgf8 morphogen gradient forms by a source-sink mechanism with freely diffusing molecules. Nature 461, 533–536.

Zacharias, D. A., Violin, J. D., Newton, A. C., and Tsien, R. Y. (2002). Partitioning of Lipid-Modified Monomeric GFPs into Membrane Microdomains of Live Cells. Science 296, 913–916.

Zecca, M., Basler, K., and Struhl, G. (1996). Direct and long-range action of a wingless morphogen gradient. Cell 87, 833–844.

Zhou, S., Lo, W. C., Suhalim, J. L., Digman, M. A., Gratton, E., Nie, Q., and Lander, A. D. (2012). Free extracellular diffusion creates the Dpp morphogen gradient of the Drosophila wing disc. Curr. Biol. 22, 668–675.

Zhu, A. J., and Scott, M. P. (2004). Incredible journey: how do developmental signals travel through tissue? Genes Dev. 18, 2985–2997.

